# A vagal-to-sympathetic reflex controls lung vascular leak

**DOI:** 10.64898/2026.10.22.761052

**Authors:** Emily K. Robinson, Emma M. Ibanez, Blanca Santibanez Ocampo, Juan Li, Nupur Ballal, Joelle M. Miller, Kaustubh Shivdikar, Benjamin D. Umans, Sara L. Prescott

## Abstract

Dynamic regulation of the vascular barrier is required for an effective inflammatory response, allowing delivery of plasma proteins and immune cells to sites of insult. Although normally protective, sustained leak becomes costly in organs like the lung, where swelling stiffens tissue and impairs breathing, yet many upstream regulators remain unidentified. Using a chemogenetic screen of vagal afferent subtypes, we identify a neural reflex that drives lung vascular leak. The reflex originates in an orphan population of neurons expressing *Cysltr2* that innervates alveolar capillary beds and senses both mechanical and inflammatory stimuli, fitting classic definitions of pulmonary “J” receptors, an afferent subtype described anatomically since the 1950s whose function has remained speculative. Activating CYSLTR2 neurons increases lung vascular leak within minutes and causes perivascular edema and respiratory dysfunction, whereas silencing them during inhaled allergen challenge reduces leak and improves lung mechanics. Rather than acting locally through antidromic peptide release, this reflex engages brainstem relays and sympathetic efferents to broadcast its effects throughout lung vasculature. Vagal control of the vascular barrier thus forms an axis of organ defense that can go awry in disease, revealing new ways to regulate organ fluid balance upstream of the endothelium.

## Introduction

The lungs are a uniquely vulnerable interface between the body and the environment, requiring tiers of neurally-coordinated defense. The vagus nerve orchestrates much of this surveillance, densely innervating the airways and other organs to provide moment-by-moment feedback to the brain^1,2^. Vagal neurons evoke protective reflexes including cough and gasp^3,4^, changes in breathing^5–7^, sickness behavior^8^, secretion^9–11^ and immune activation^9–11^. Underlying this functional breadth is a diversity of afferent neural pathways^3,12–16^, yet even in an organ as important as the lungs, many afferent subtypes have no assigned function, raising the possibility that entire classes of vagal reflexes may have eluded description.

One element of lung defense in which the vagal inputs are poorly defined concerns the regulation of the vascular barrier. Regulated increases in endothelial permeability are an important arm of innate immunity across many tissues, facilitating the delivery of immune cells, antibodies, and plasma proteins to sites of insult^17–20^. In the skin, sensory neurons promote vascular leak through the antidromic release of tachykinins, a phenomenon known as neurogenic inflammation^21,22^. Electrical, pharmacological, and vagotomy studies suggest vagal neurons may control vascular leak in the lungs as well^23–25^, but the precise afferent or effector mechanisms involved have not been identified. Furthermore, though some subsets of vagal lung afferents express tachykinins, the sparsity of substance P-positive neurons near sites of barrier disruption^24^ and the failure of tachykinin receptor antagonists in human clinical trials cast doubt on the relevance of a local peptidergic mechanism in the lungs^26^. Vagal neurons may therefore modulate vascular barriers by mechanisms distinct from those employed in the skin.

Although vascular permeability is understood to facilitate the resolution of acute injury, excessive or sustained leak produces edema that compromises respiratory function. The nature of that compromise, and its relative participation in different diseases contexts, depends on which vascular compartment is breached. For example, disruption of the alveolar-capillary barrier causes flooding of the alveolar septa and air spaces that is central to the pathogenesis of pneumonia^27,28^ and acute respiratory distress syndrome (ARDS) ^29,30^, the leading cause of COVID-19 mortality. Alternatively, excessive leak from the bronchial microvasculature causes swelling of the airway wall, exacerbating the airflow restriction and remodeling seen in asthma^31,33^. Finally, leak from post-capillary venules causes peribronchovascular cuffing, fluid buildup in the interstitial zones around large blood vessels, disrupting lung mechanics in conditions like cardiogenic or high-altitude pulmonary edema^32–35^ and vasculitis^32–35^. Altered vagal signaling is documented in many of these conditions^32–35^, although its significance is unclear. Despite decades of work on endothelial barrier regulation, edema cannot yet be treated at its source^36,37^, underscoring a need to identify additional upstream regulators of permeability.

Here we identify a previously uncharacterized vagal sensory population that controls lung vascular leak. These neurons are marked by expression of the cysteinyl leukotriene receptor *Cysltr2*, innervate the alveolar capillary bed, and respond to mechanical and inflammatory stimuli--resembling the pulmonary J (“juxtacapillary”) receptors described by Paintal in the mid-20^th^ century^38,39^. Recurrently activating CYSLTR2 neurons drives fluid accumulation around the airways and vessels and impairs lung mechanics, whereas silencing them during allergic challenge reduces perivascular cuffing and preserves lung function. Unlike leak in the skin, vagally evoked edema requires brainstem relays and sympathetic efferents rather than antidromic release. Together, these findings establish a previously unrecognized vagal-to-sympathetic reflex that has wide implications for both lung health and disease.

## Results

### A chemogenetic screen identifies a vagal subtype that persistently alters lung mechanics

Many respiratory diseases are characterized by tissue changes that persistently alter airway mechanics. While most prior studies of neural influence on lung function have focused on immediate reflex effects, we sought to determine whether persistent or mistuned vagal signaling is sufficient to drive lasting features of airway dysfunction. To assess this possibility, we gained genetic access to lung-innervating vagal neurons using a panel of subtype-restricted *Cre* driver lines. Lung afferents have been identified through both tracing^3,4,15,40^ and lung retrograde labeling studies^13,16^, encompassing at least 10 vagal populations of both known and unknown functions (Fig. 1A, S1A-B). Our *Cre* panel included predicted chemosensory and mechanosensory subtypes, including jugular (*Npy1r-Cre*^3^) and nodose C-fiber (*Vip-Cre*^41^ and *Cysltr2-Cre)* populations, and large-diameter neurons (*Gabra1-Cre*^3,40^*, Calb1-Cre*^3,15^) which innervate pharyngeal and trachealis terminals. Because *Calb1-Cre* exhibits lineage artifacts, we substituted *Kcng4*-*Cre*, which targets a jugular *Calb1^+^* subset that recapitulates tracheal chandelier-shaped terminals upon vagal tracing (Fig. S1B). *Vglut2-Cre*, which broadly accesses all vagal afferents^5^, as well as *Piezo2-Cre,* a marker for vagal mechanoreceptors^42^, were included as controls. Together, our panel covers the majority of reported lung chemo- and mechanosensory neurons.

**Fig. 1.**
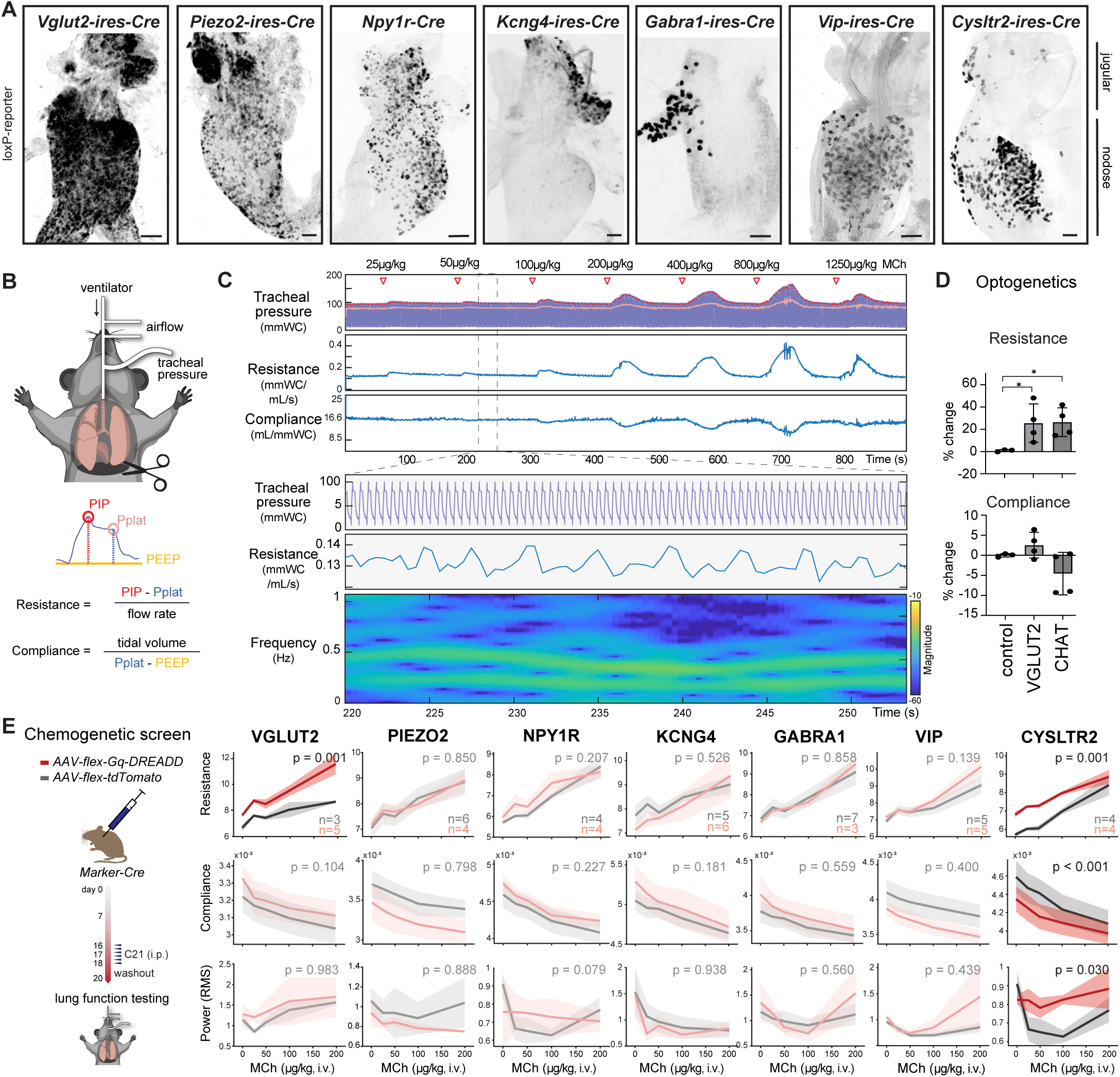
A chemogenetic screen identifies a lung afferent subtype that persistently alters airway mechanics. **(A)** Vagal ganglia from mice carrying *Cre* alleles and *Cre*-dependent reporters marking seven lung-innervating vagal sensory populations (*loxP-ChR2-eYFP* for *Vglut2-ires-Cre*, *Gabra1-ires-Cre, Vip-ires-Cre* and *Cysltr2-ires-Cre*; *loxP-DTR* for *Piezo2*-*ires-Cre*, *Kcng4*-*Cre*, and *Npy1r*-*Cre*). Scale bars, 100 μm. **(B)** Lung function testing paradigm and derived parameters. **(C)** Top: representative tracheal pressure trace (mmWC; 10 mmWC = 1 cmH₂O) with peak inspiratory (red) and plateau (pink) pressures marked during methacholine (MCh) challenge in a wildtype animal, above the derived resistance (mmWC/mL/s) and compliance (mL/mmWC) measurements. Bottom: expanded view of breath-to-breath resistance oscillations and a spectrogram of detrended resistance showing spectral power (dB) versus frequency (Hz) and time. Color axis represents the magnitude in dB relative to 1 mmWC·min/mL. **(D)** Change in resistance and compliance during optogenetic stimulation of the vagosympathetic trunk in mice expressing *ChR2* from the indicated *Cre* drivers. n = 3–4 mice per line; two-tailed unpaired t test; *P < 0.05, **P < 0.01. **(E)** Left: design of the DREADD-based chemogenetic screen. Right: resistance (AUC), compliance (median), and wave power (AUC of RMS spectral power) over the 60 s after each MCh dose (i.v.; 25, 50, 100, 200 μg/kg) in mice injected with *AAV-Flex-tdTomato* (control, gray) or *AAV-hSyn-DIO-hM3D(Gq)-mCherry* (red). See supplementary materials for additional information on lung function metrics. P-values were calculated using an omnibus model with dose as a categorical factor, group × dose interaction reported; Data are means ± SEM throughout.

To screen for vagal pathways that shape lung health, we turned to lung function testing. Lung function tests report on the mechanics of breathing, principally resistance (the pressure needed to drive air through the airways, which reflects airway narrowing), compliance (the ease with which the lungs expand for a given pressure, which reflects tissue stiffness), and their temporal fluctuation (the breath-to-breath variability of these values over time that carries independent prognostic information in airway disease^43–45)^. Since most commercial systems for mouse lung function testing are incompatible with intravital vagal manipulation, we developed a custom platform that extracts these parameters in real time from a recurrent breath-hold maneuver in anesthetized animals. The design offers two advantages: every breath cycle is resolved rather than sampled intermittently, so we measure fluctuation directly, and the neck remains exposed for vagal access (Fig. 1B; details in supplementary methods). To illustrate the sensitivity of the system, we performed methacholine bronchoprovocation testing, which showed reversible and dose-dependent changes in resistance and compliance (Fig. 1C, top) and recapitulated known sex and age effects (Fig. S1C). A moving window of variation revealed slow, rhythmic oscillations in baseline airway resistance (∼0.5 Hz, Fig. 1C, bottom), reminiscent of the airway peristalsis reported in neonatal and juvenile mice and in other species^46–48^. Wave power of these oscillations decreased transiently after each methacholine dose, as expected for a slow, coordinated contraction. To illustrate our platform’s compatibility with vagal manipulations, we performed optogenetic activation of all vagal sensory (*Vglut2-Cre;loxP-ChR2*) or motor (*Chat-Cre;loxP-ChR2*) neurons, which immediately increased airway resistance without altering compliance (Fig. 1D). These effects resolved rapidly after light stimulus cessation. Thus, our platform provides a readout to link genetically-defined neural pathways to both acute and persistent changes in lung mechanics *in vivo*.

These findings confirm that lung mechanical changes are largely reversible following transient vagal activation. We next performed a chemogenetic screen across our *Cre* line panel to constitutively activate vagal pathways for several days, mimicking the recurrent activation often seen with naturalistic stimuli, then incorporated a washout period (Fig. 1E) to isolate effects that persist beyond neuronal activation. This strategy was designed to remove acute reflex components and instead unmask previously-hidden slower effects with potential disease relevance, such as reflex circuit plasticity, lung structural remodeling, immune sequelae or other lasting tissue changes. For each *Cre* line, we delivered a *Cre*-dependent excitatory DREADD^49^ (*AAV-Gαq-DREADD)* to the vagal ganglia by AAV injection. After 3 weeks of recovery, we administered the synthetic ligand compound 21 (C21)^50^ twice daily for 3 days by intraperitoneal injection followed by a 2-day washout before terminal lung function testing. Activation of all vagal sensory neurons (*Vglut2-Cre*) led to an increase in airway resistance, demonstrating that repeated vagal activation drives lasting mechanical changes in the absence of lung injury or a primed allergic response (Fig. 1E, left). Activation of PIEZO2-, NPY1R-, KCNG4-, VIP-, or GABRA1-defined vagal subpopulations produced no lasting effects. Strikingly, activation of CYSLTR2 neurons produced a sustained elevation in resistance, decrease in compliance, and heightened airway lability (Fig. 1E, right). Because CYSLTR2 neurons are a population of unknown function, we sought to further characterize their role in lung physiology.

### CYSLTR2 neurons are alveolar “juxtacapillary” (J) receptors

We first examined CYSLTR2 neuron identity. Sensory neurons have historically been classified in part by their end terminal location and morphology, so we mapped CYSLTR2 neurons to their peripheral targets using genetically-guided viral tracing. CYSLTR2 neuron innervation outside the lung was sparse, limited to the heart and gut mucosa and consistent with *Cre* expression in other rare sensory subtypes (Fig. S2A). Because the *Cysltr2-Cre* line had not previously been validated in vagal neurons, and to assess potential lineage effects, we also performed two-color RNA *in situ* hybridization against *Cysltr2* and *ChR2* transcripts in the vagal ganglia of *Cysltr2-Cre;loxP-ChR2* mice. Few cells expressed *ChR2* without detectable *Cysltr2* expression (∼8% of all *Chr2*-expressing cells, Fig. S2C,D), indicating limited off-target effects.

Within the lung, the majority of traced CYSLTR2 fibers extended along major bronchioles before diverting into the alveolar compartment (Fig. 2A). Individual fibers then branched and encircled one or multiple alveoli before terminating within a CD31+ capillary bed (Fig. 2B, Movies 1 and 2). At higher magnification, interseptal capillary terminals took two forms: flattened elaborate arbors (“tree branch”), or small beaded (“rosebud”) endings. Both intercalated near capillary junctions, with occasional terminals also near neuroepithelial bodies (Fig. S2B). This terminal position within the alveolar wall is the defining anatomical criterion of pulmonary J (“juxtacapillary”) receptors.

**Fig. 2.**
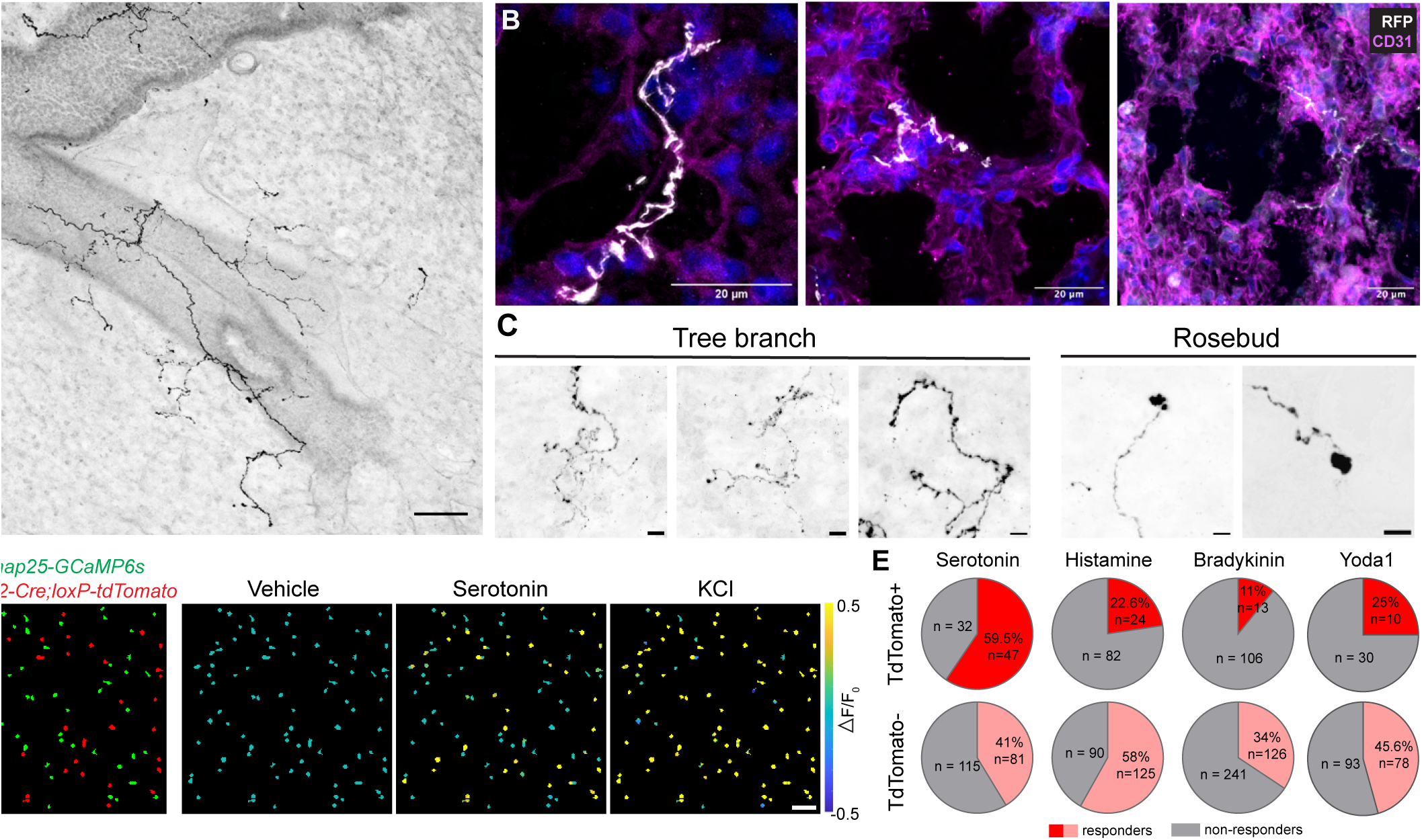
CYSLTR2 neurons are pulmonary J receptors. **(A)** Labeled terminals in a 300-μm lung slice from *Cysltr2-ires-Cre* mice injected in the nodose/jugular/petrosal (NJP) ganglia with *AAV-Flex-tdTomato*. Scale bar, 100 μm. **(B)** *Cysltr2*⁺ terminals (white) adjacent to alveolar structures labeled by CD31 (magenta), with DAPI nuclear counterstain in blue. Scale bar, 20 μm. **(C)** Example morphologies of *Cysltr2⁺* terminals. Scale bars, 10 μm. **(D)** Calcium imaging of dissociated vagal neurons from *Cysltr2-Cre;loxP-tdTomato;Snap25-GCaMP6s* mice during sequential stimulus perfusion. Left: Snapshot showing all vagal neurons expressing *Snap25-GCaMP6s* with a subset marked by *tdTomato*; scale bar, 100 μm. Right: Heatmap overlay showing ΔF/F₀ per cell following vehicle (saline), serotonin (10 µM), and KCl (50mM) stimulus. **(E)** Fraction of *tdTomato*⁺ and *tdTomato*⁻ neurons responding to each stimulus.

Classically, pulmonary J receptors were defined as slowly-conducting neurons with dual sensitivity to specific chemicals (serotonin) and mechanical stimuli (raised capillary pressure, lung hyperinflation)^51^. *Cysltr2* marks a population of C-fiber lung afferents of nodose origin (*Phox2b^+^*, *Nefh-*low, *Scn10a^+^, Htr3a^+^, Piezo1^+^*) that potentially fits this profile. To ask if CYSLTR2 neurons share these sensory properties, we tested the responses of CYSLTR2 neurons to classic pulmonary J receptor stimuli *in vitro*. We dissociated vagal neurons from *Snap25-GCaMP6s;Cysltr2-Cre;loxP-tdTomato* mice and recorded calcium responses in CYSLTR2^+^ and CYSLTR2^−^ neurons in parallel during *ex vivo* perfusion (Fig. 2D, 2E). CYSLTR2^+^ neurons had enriched responses to serotonin, the prototypical J receptor agonist, compared to CYSLTR2^−^ neurons (59.5% versus 41%, respectively). In contrast, responses to bradykinin and histamine, which act on other lung C-fiber classes but spare J receptors, were depleted. Given the expression of *Cysltr2*, we also tested their responsiveness to cysteinyl leukotrienes C4 and D4^52,53^. These did not elicit detectable calcium responses in any neurons tested, suggesting a modulatory rather than directly excitatory function (Fig. S2E).

Another criterion for J receptors is their sensitivity to mechanical cues. Interestingly, while lung-innervating CYSLTR2 neurons do not express mechanosensitive ion channel *Piezo2* (a marker for most visceral mechanoreceptors), they express high levels of *Piezo1*. We therefore tested calcium responses to the *Piezo1* agonist Yoda1. We observed Yoda1-induced activation of a subset of CYSLTR2 neurons, consistent with the *Cysltr2-Cre* expression encompassing the *Piezo1*^+^ lung afferent subtype; Yoda1 responses were observed in CYSLTR2^-^ neurons as well, as expected given the presence of *Piezo2^+^Piezo1^+^Cysltr2^-^*baroreceptors that innervate the aortic arch^51^. Thus, CYSLTR2 neurons fit the anatomical and sensory criteria of pulmonary J receptors, providing a genetic handle to interrogate their function.

### CYSLTR2 neurons cause tissue congestion by inducing vascular leak

We therefore sought to determine how their persistent activation impairs lung function. We first histologically examined the lungs of the chemogenetically-activated *Cysltr2-Cre* mice to look for structural clues. Their lungs showed no evidence of smooth muscle hypertrophy, fibrosis, goblet cell hyperplasia, or excessive immune infiltration (Fig. S3A). The only structural correlate was perivascular cuffing, a sign of fluid overload in the interstitial spaces surrounding the large blood vessels that often precedes more severe alveolar edema (Fig. 3A, 3B). Interstitial fluid accumulation around the bronchovascular bundle can disrupt lung function by compressing airway walls and mechanically uncoupling them from surrounding parenchyma--findings that could explain both the increased resistance and decreased compliance seen in our lung function tests^54^. Because cuffing is typically considered a leading rather than lagging sign of abnormal fluid homeostasis, we next turned to more acute readouts of edema. We therefore applied vagal optogenetics and measured the post-mortem gravimetric lung wet-to-dry (W/D) ratio, a standard metric of edema^55,56^. Even just a brief 10-minute stimulation (20Hz, 5ms) of CYSLTR2 neurons raised the wet-to-dry (W/D) ratio by ∼45% compared to sham-stimulated animals, exceeding the effect of intratracheal histamine over the same interval (∼23% W/D ratio increase) (Fig. 3C). This rapid rise indicates extravascular fluid accumulation.

**Fig. 3.**
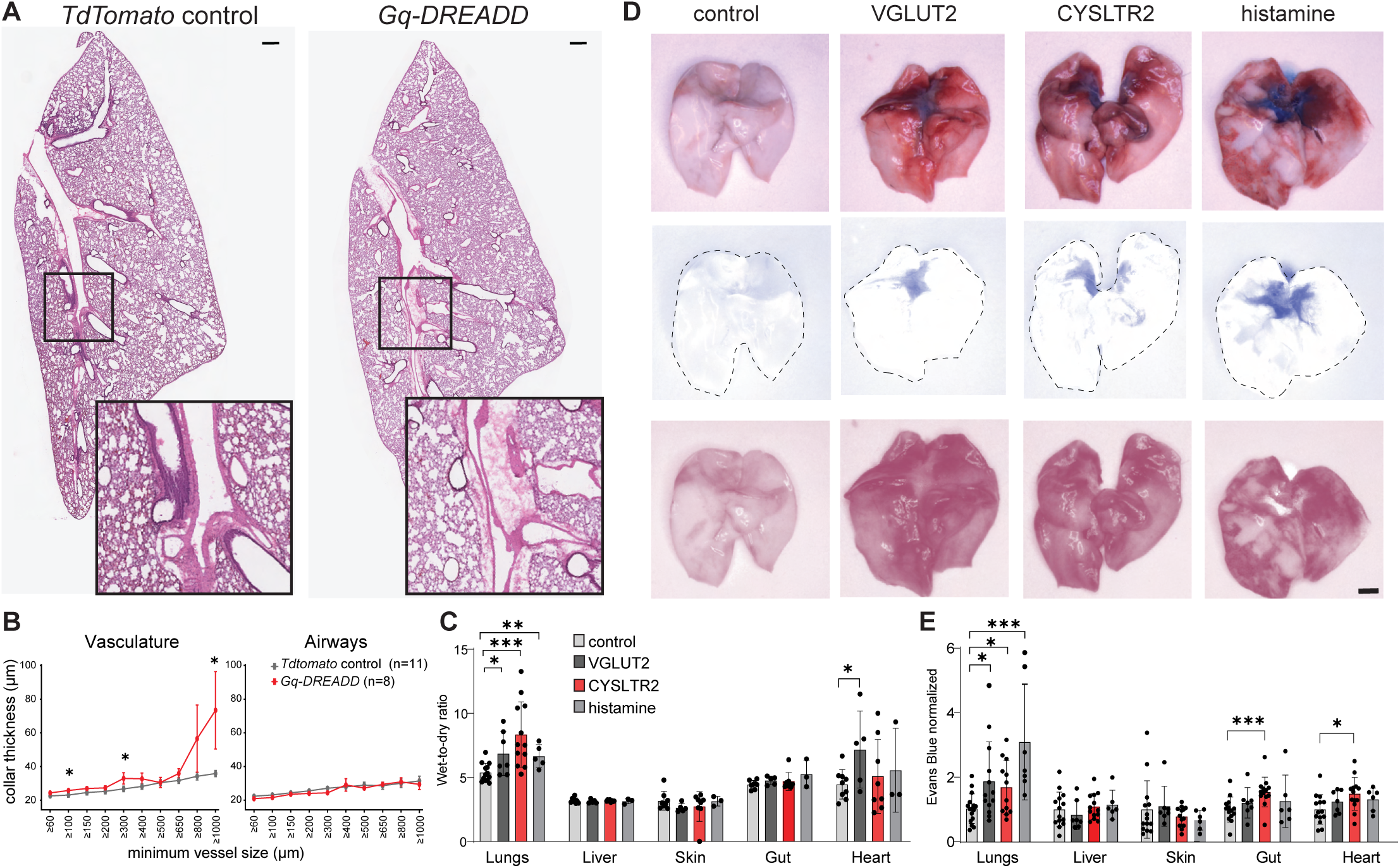
CYSLTR2 neurons induce vascular leak. **(A)** Hematoxylin and eosin (H&E)–stained lungs from *Cysltr2-ires-Cre* mice injected in the vagal ganglia with *AAV-Flex-tdTomato* (control) or *AAV-hSyn-DIO-hM3D(Gq)-mCherry* (Gq-DREADD). Bottom: higher magnification of major pulmonary vessels showing perivascular cuffing. Scale bar, 1 mm. **(B)** Collar thickness (μm) surrounding pulmonary blood vessels (left) or airways (right), binned by minimum vessel diameter, in *tdTomato* control (gray) and *Gq-DREADD* (red) mice. n = 8-11 mice per group; two-tailed unpaired Welch’s t test per bin. **(C)** Wet-to-dry ratios of lung, liver, skin, gut, and heart after optogenetic stimulation (50Hz, 5ms pulses, 10s on-20s off for 10 minutes) in *Cre*-negative *loxP-ChR2* controls (light gray), *Vglut2-Cre;loxP-ChR2* (dark gray), and *Cysltr2-Cre;loxP-ChR2* (red) mice, and in mice given endotracheal histamine (0.15 mg, medium gray). n = 3–14 mice per group; two-tailed unpaired t test. **(D)** Representative whole lung lobes (top) with Evans blue (middle) and red (bottom) channels separated by color deconvolution (see supplementary materials). Scale bar, 2 mm. **(E)** Evans blue absorbance normalized to the mean absorbance of control lobes; groups and colors and stim conditions as in (C). n = 7–14 mice per group; two-tailed unpaired t test. Asterisks as in Fig. 1D; data are means ± SEM.

Fluid accumulation in the lungs can have various causes. Since peribronchovascular cuffing is frequently seen in cardiogenic (heart-related) pulmonary edema, we asked if this fluid buildup was secondary to impaired cardiac function. Unlike VGLUT2 neuron stimulation, which evoked immediate tachypnea and bradycardia, transient (10-second) activation of CYSLTR2 neurons had no immediate effects on heart rate or breathing, suggesting that the edema is non-cardiogenic and precedes later respiratory dysfunction (Fig S3B, S3C). Another possibility is that CYSLTR2 neurons evoke edema by increasing endothelial permeability. We therefore assessed vascular barrier integrity using an Evans blue assay, a blue dye that binds albumin and allows direct quantification of plasma leak. A sustained 10-minute optogenetic activation of either VGLUT2 or CYSLTR2 neurons induced profound endothelial barrier disruption. After neuron activation, lungs were visibly bluer and had greater tissue congestion than control-stimulated lungs, similar to the lungs of histamine-treated mice (Fig. 3D), indicating accumulation of plasma albumin. Lung dye extraction confirmed ∼54% increase in extravasation following CYSLTR2 neuron activation compared to sham controls (Fig. 3E). Thus, CYSLTR2 neuron activation rapidly disrupts the vascular barrier, causing peribronchovascular edema that impairs lung function.

### CYSLTR2 neurons are engaged during an inhaled ovalbumin challenge

Having found that activating CYSLTR2 neurons is sufficient to drive lung peribronchovascular leak, we asked whether these neurons participate in tissue congestion caused by an inhaled challenge. Cuffing (sometimes described as a ‘donut sign’) is a common radiologic sign in airway allergy^57,58^. Therefore, we turned to an inhaled ovalbumin (OVA) model for its simplicity, high degree of vascular involvement^57,58^, and compatibility with genetic manipulation. Mice were sensitized with OVA in aluminum hydroxide (alum) on days 0 and 7, then challenged with inhaled OVA on three consecutive days; controls received alum alone followed by inhaled saline (PBS). By day 20, OVA-challenged animals exhibited increased serum OVA-specific IgE levels, immune infiltration, and tissue remodeling, confirming an allergic response (Figs. 4A, S5A, S5B, S5C, S5F). As expected, the lungs of OVA-treated mice showed pronounced perivascular cuffing, with many vessels surrounded by an edematous zone with immune infiltration (Fig. 4A, S5E). This was associated with lung function impairment, marked by increased resistance and reduced compliance—collectively a stiffer, flow-limited phenotype (Fig. 4B, S5D). Airway hyperresponsiveness was not observed, as commonly reported for allergen models in the C57BL/6 background^59^.

**Fig. 4.**
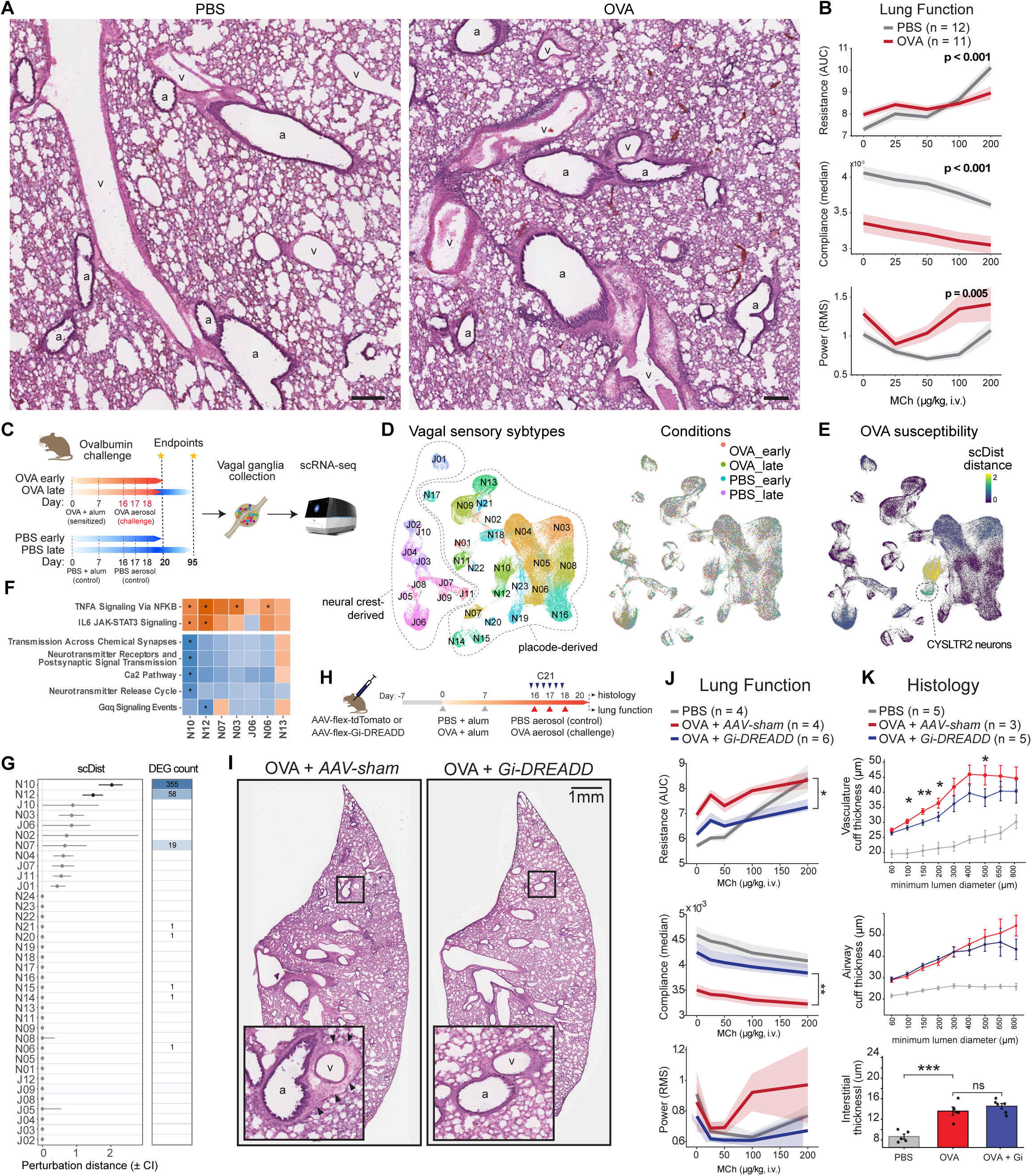
CYSLTR2 neurons respond to airway allergens and mediate allergen-evoked vascular leak in the lungs. **(A)** H&E-stained lung sections from PBS- and ovalbumin (OVA)–treated mice showing perivascular cuffing; a, airway; v, vessel. Scale bar, 250 μm. **(B)** Resistance (AUC), compliance (median), and wave power during MCh challenge (i.v.; 25, 50, 100, 200 μg/kg) in PBS-treated (gray, n = 12) and OVA-treated (red, n = 11) mice. P-values reflect omnibus model as in Fig. 1E. **(C)** Schematic of single-cell transcriptomic analysis, showing collection endpoints. **(D)** Integrated UMAP showing vagal sensory neuron subtypes, pooled across all PBS- and OVA-treated mice at early and late endpoints (20 mice per condition; 129,810 cells total). Left: jugular (J1–J10) and nodose/petrosal (N1–N23) clusters are annotated. Right: same UMAP pseudocolored by condition. **(E)** Same UMAP pseudocolored by scDist perturbation distance. **(F)** Selected enriched gene sets of OVA-altered genes in clusters N10, N12, N03, J06, N06, and N13, calculated with fgsea; colored by normalized enrichment score, asterisks indicate significant enrichment (q<0.05). **(G)** scDist perturbation distance (±95% CI) and differentially expressed gene (DEG) count per nodose/jugular population. **(H)** Schematic of timeline following intranodose injection of *AAV-Flex-tdTomato* or *AAV-hSyn-DIO-hM3D(Gi)-mCherry*, with the PBS or OVA sensitization-and-challenge. Blue arrows indicate C21 dosing (i.p., 3 mg/kg, twice daily), red arrows indicate OVA aerosol during challenge. **(I)** H&E-stained lung sections from OVA-treated *Cysltr2-ires-Cre* mice injected in the nodose ganglia with *AAV-Flex-tdTomato* (OVA + *AAV-sham*) or *AAV-hSyn-DIO-hM3D(Gi)-mCherry* (OVA + *Gi-DREADD*), with inset showing higher magnification of major pulmonary vessels (v) and airways (a); black arrows indicate cuffing. Scale bar, 1 mm. **(J)** Resistance (AUC), compliance (median), and wave power during MCh challenge (doses as in (B)) in mice treated with PBS + *AAV-Flex-tdTomato* (gray, n = 4), OVA + *AAV-Flex-tdTomato* (red, n = 4), and OVA + *AAV-hSyn-DIO-hM3D(Gi)-mCherry* (blue, n = 6). P-value reflects omnibus model, as in Fig. 1E; only the p-value for OVA + AAV-sham versus OVA + AAV-Gi-DREADD is shown. For PBS versus OVA, PBS versus OVA + Gi, and OVA versus OVA + Gi, respectively: resistance, P < 0.001, P < 0.001, P = 0.019; compliance, P < 0.001, P = 0.009, P = 0.002; wave power, P = 0.314, P = 0.439, P = 0.070. **(K)** Different histological metrics were quantified in mice treated with PBS (gray, n = 5), OVA (red, n = 3 cuff, n = 5 interstitial), and OVA + Gi (blue, n = 5 cuff, n = 6 interstitial). Top: perivascular cuff thickness, binned by minimum vessel diameter. Middle: peribronchial cuff thickness, binned by minimum airway diameter. Bottom: interstitial thickness. Welch’s unpaired t test; asterisks denote OVA versus OVA + Gi; PBS differed from both OVA and OVA + Gi at P < 0.0001 in all bins. Asterisks as in Fig. 1D; data are means ± SEM.

To ask whether CYSLTR2 neurons participate in OVA-induced vascular edema, we turned to single-cell RNA sequencing. Since activated vagal neurons are not effectively labeled by c-FOS, we reasoned alternatively we could use changes in gene expression to predict functionally-engaged subtypes. We harvested vagal ganglia from mice subjected to OVA or PBS challenge, collected at the peak of airway inflammation (day 20), or 8 weeks following the last OVA or PBS challenge (Fig. 4C) to assess signs of long-lasting transcriptional memory. In total we captured ∼323K cells across the four conditions, including vagal neurons, sympathetic neurons from the nearby superior cervical ganglion, satellite glia, Schwann cells, immune cells, endothelial cells, and fibroblasts (Fig. S4A; n=5 male mice per replicate, four replicates per condition). Post-filtering we recovered 129,810 sensory neurons which clustered into 36 vagal and glossopharyngeal populations, which we enumerated as J1-10 and N1-26, for “jugular” and “nodose/petrosal” (Figure 4D). Each subtype mapped cleanly to previously published vagal/glossopharyngeal atlas clusters (Fig S4B). All subtypes were recovered in similar proportions across conditions, with no evidence of cluster depletion or transdifferentiation following OVA treatment, indicating that cell-type identity was preserved (Fig. 4D, S4C). While statistically-significant gene expression changes were observed in a subset of clusters during the peak of airway inflammation (day 20), nearly all gene expression changes reverted by 8 weeks post-recovery despite lasting lung function deficits (Fig. S4D, S4E). This demonstrates that ovalbumin-induced airway challenge elicits robust and reversible transcriptional responses in a subset of vagal sensory neurons.

To prioritize vagal subtypes with the strongest responses to inhaled OVA, we next ranked clusters by (i) per-cluster counts of differentially expressed genes in the acute timepoint, and (ii) scDist^60^, a mixed-effects framework for quantifying transcriptional shifts that accounts for inter-animal variability and differences in cluster size. Each method independently converged on two nodose populations with the largest transcriptional responses—N10 and N12 (Fig. 4G). Both subtypes preferentially innervate the lungs in retrograde labeling studies^14,17^, supporting the biological relevance of our analysis. The strongest responder, N10, corresponds to a lung afferent subtype marked by *Vip* that was recently shown to drive allergen-induced airway hyperresponsiveness^41^. The second class, N12, corresponds to the lung afferent subtype targeted by *Cysltr2-Cre*, confirming their functional engagement during OVA-induced airway inflammation. The major markers and receptor profiles of both populations remained largely unchanged by the OVA challenge; however, gene set enrichment analysis (GSEA) predicted changes in genes related to neuronal plasticity, TNFα–NFκB, and JAK–STAT3 signaling pathways (Fig. 4F, Table S3). Both N10 and N12 neurons strongly express the TNFα receptor *Tnfrsf1a* and Janus kinase (JAK) family machinery *Jak1/Stat3/Stat6* (Fig S4F), consistent with the known roles of TNFα and JAK-STAT signaling in airway allergy and neuronal sensitization^1,61^, providing a cell-intrinsic basis for their favored engagement.

The observation that only 2 out of 36 vagal populations respond to OVA challenge supports the labeled-line model of viscerosensory processing, and motivated closer comparison of their molecular identities (Fig. 4E). Both N10 and N12 are placode-derived (*Phox2b*^+^) and express a broad repertoire of receptors for inflammatory mediators (*Tnfrsf1a*, *Il13ra1*, *Nod1*, *Ednra*, *S1pr3*, *Ptger2/4*). They diverge however in core identity. N10 neurons are prototypical C-fibers, marked by TRP channels (*Trpv1*/*Trpa1*), neuropeptides (*Calca/Calcb, Vip*), and sodium channels (*Scn10a*, *Scn11a*). N12 neurons, by contrast, lack TRP channels and sensory neuropeptides (*Tac1/2/3*, *Calca/Calcb*, *Vip*) and instead express a combined mechano-/chemosensory module (*Adgrl2*, *P2rx3*, *P2ry1*, *Htr3a/b*, *Scn7a*, *Scn10a*), including *Piezo1* but not *Piezo2*, as described above (Fig. S4G). The observation that N12 neurons lack *Trpv1* expression was notable given the widespread use of capsaicin to identify vagal C-fibers, despite reports of capsaicin-insensitive C-fibers *in vivo*^62^. We therefore tested capsaicin responsiveness of CYSLTR2 neurons using *ex vivo* calcium imaging and found that all Yoda1-responsive *tdTomato*-marked neurons were universally capsaicin-insensitive (Fig S4H, S4I). Together, this identifies two distinct pathways that are preferentially engaged during allergic airway inflammation, including one comprised of a *Cysltr2^+^Piezo1^+^Trpv1^-^*neuronal subtype.

### Silencing CYSLTR2 neurons mitigates edema and restores lung function during OVA challenge

Given that more than one vagal pathway was activated by OVA, we asked whether silencing CYSLTR2 neurons alone mitigates OVA-induced pathology. We injected the vagal ganglia of *Cysltr2-Cre* mice with virus carrying a *Cre*-dependent inhibitory DREADD (*AAV-Gi-DREADD-mCherry).* We inhibited CYSLTR2 neurons with C21 (twice daily, i.p.) throughout the OVA aerosol challenge, after immune memory has already formed (Fig 4H). CYSLTR2-silenced mice had similar levels of OVA-specific IgE serum levels, goblet cell hyperplasia, and total immune cell burden, indicating that their allergic response was not impaired (Fig. S5A-C). However, compared to OVA-treated animals injected with an *AAV-mCherry* control virus, the lung function of CYSLTR2-silenced mice was significantly improved, showing reduced airway resistance, instability, and elastance (Fig. 4J, S5D). Evaluation of lung histopathology also showed significantly reduced cuffing, particularly around large-diameter vessels, but unchanged airway wall and alveolar interseptal thickness, suggesting a regionalized effect on post-capillary venule permeability (Fig 4I, 4K, S5E-F). Silencing of CYSLTR2 neurons therefore prevents tissue congestion.

### The vagal vascular leak reflex is mediated through central and not peripheral pathways

Neurogenic vascular leak can be mediated through either local (antidromic) or central mechanisms. In the skin, somatosensory neurons drive leak by releasing tachykinins onto nearby endothelial and immune cells, leading to the characteristic ‘wheal-and-flare’ of allergic skin reactions. However, CYSLTR2 neurons lack *Tac1/2/4* (Fig. S4G) and are located at a distance from the site of fluid accumulation, arguing against a local antidromic mechanism. Additionally, substance P-positive neurons are sparse in humans, and not found the vicinity of leaky vessels^24,26^, casting doubt on the importance of this pathway in the lung. We therefore considered a model in which vagally-evoked leak involves central reflex circuits.

To distinguish peripheral (antidromic) from central (orthodromic) mediation, we combined targeted chemogenetic activation with nerve cuts. *Cysltr2-Cre* mice were injected unilaterally with an excitatory DREADD (*AAV-flex-Gq-DREADD-mCherry*), and one vagal trunk was transected either ipsilaterally below the vagal ganglion, eliminating peripheral axons while preserving central connections, or contralaterally, preserving both (Fig, 5A). We did not cut above the ganglion because the vagal rootlets are surgically inaccessible inside the murine skull. After 2-week recovery, we unilaterally activated CYSLTR2 neurons with intraperitoneal C21 and measured lung function. Unilateral activation without a nerve cut, or with a contralateral nerve cut, showed lung function impairment comparable to uncut bilateral CYSLTR2 activation (Fig. 5A,S6A). Ipsilateral nerve cuts also did not abolish the DREADD-induced changes in resistance and compliance, indicating that antidromic signaling was not required for the effect (Fig. 5A). Thus, the effects of CYSLTR2 neurons on lung function are mediated at least partly through central relays rather than by direct action of sensory neurons on lung tissue.

**Fig. 5.**
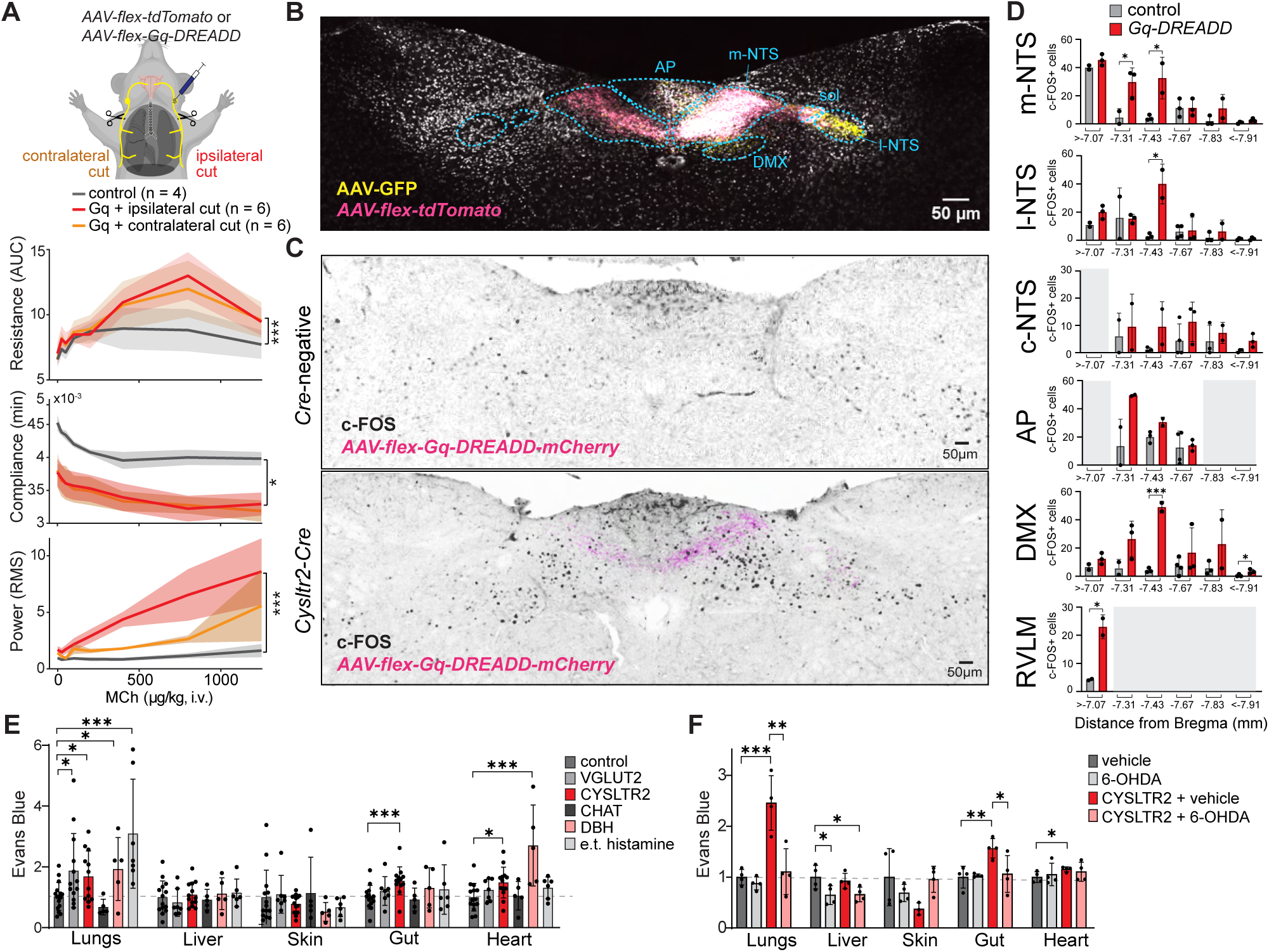
Pulmonary vascular leak is mediated by a central vagal-sympathetic reflex circuit. **(A)** Top: Schematic of unilateral CYSLTR2 neuron activation combined with ipsilateral or contralateral vagotomy. Bottom: resistance (AUC), compliance (median), and wave power (AUC RM during MCh challenge (i.v.; 25, 50, 100, 200, 400, 800, 1250 μg/kg) in *Cysltr2-ires-Cre* mice injected unilaterally with *AAV-Flex-tdTomato* (gray, n = 4), unilaterally with *AAV-hSyn-DIO-hM3D(Gq)-mCherry* with ipsilateral vagotomy (red, n = 6), or unilaterally with *AAV-hSyn-DIO-hM3D(Gq)-mCherry* with contralateral vagotomy (orange, n = 6). P-values reflect omnibus model, as in Fig. 1E. For control versus Gq-ipsilateral, control versus Gq-contralateral, and Gq-ipsilateral versus Gq-contralateral, respectively: resistance, P = 0.002, P = 0.199, P = 0.948; compliance, P = 0.044, P = 0.076, P = 0.907; wave power, P < 0.001, P = 0.003, P = 0.336. Asterisks denote control versus Gq-ipsilateral. **(B)** A brainstem section from a *Cysltr2-ires-Cre* mouse unilaterally injected with *AAV-Flex-tdTomato* and *pENN.AAV.CB7.CI.eGFP.WPRE.rBG*. Scale bars, 50 μm; bregma −7.43 mm. **(C)** c-Fos (black) and mCherry (magenta) immunostaining in *Cre*-negative and *Cysltr2*-ires-*Cre* mice injected with *AAV-hSyn-DIO-hM3D(Gq)-mCherry*, 90 minutes after i.p. C21. Scale bar, 50 μm. **(D)** c-Fos⁺ neuron counts in m-NTS, l-NTS, c-NTS, AP, DMV, and RVLM, binned by anatomical level (distance from bregma), n = 2–4 mice; two-tailed unpaired t test. **(E)** Vascular leak was quantified by Evans blue absorbance after optogenetic stimulation (5ms pulses, 10s on/20s off for 10 minutes) of *Cre*-negative controls (gray; 50Hz), *Vglut2*-*Cre* (light gray; 50Hz), *Cysltr2*-*Cre* (red; 50Hz), *Chat*-*Cre* (dark gray; 10Hz), or *Dbh-Cre* (pink; 10Hz) neurons. Values for each organ were normalized to the mean absorbance of controls. Endotracheal histamine (0.15 mg, grey) was provided as a positive control. n = 7–14 mice per group; two-tailed unpaired t test. **(F)** Evans blue absorbance normalized to the mean of vehicle control lobes in vehicle-treated (gray) and 6-OHDA–treated (light gray) mice, and in *Cysltr2-Cre;loxP-ChR2* mice given vehicle (red) or 6-OHDA (pink). All mice were optogenetically stimulated using same parameters as in (E). n = 4 mice per group; two-tailed unpaired t test. Asterisks as in Fig. 1D; data are means ± SEM. AP, area postrema; DMV, dorsal motor nucleus of the vagus; NTS, nucleus of the solitary tract (m-, medial; l-, lateral; c-, commissural); RVLM, rostral ventrolateral medulla.

Given that the reflex requires brainstem relay, we characterized the central projection patterns of CYSLTR2 neurons. Vagal ganglia of *Cysltr2-Cre* mice were injected with both a *Cre*-dependent *AAV-flex-tdTomato* and *Cre*-independent *AAV-eGFP* to visualize *Cre*-expressing neurons in the context of all types of vagal sensory fibers. While GFP^+^ fibers spanned the nucleus tractus solitarius (NTS), area postrema (AP), and paratrigeminal nucleus (Pa5), CYSLTR2 neurons arborized in an anatomically restricted manner, with highest labeling in the dorsomedial NTS, ipsilateral area postrema, and commissural regions caudal to the obex--regions associated with pulmonary C-fiber input^63,64^ (Fig. 5B, S6B). To exclude signals coming from fibers of passage, we combined chemogenetic activation of CYSLTR2 neurons with c-FOS mapping. Activation increased c-FOS within 90 minutes in the dorsomedial NTS and area postrema, with strongest labeling near −7.43 mm from bregma. Because c-FOS mapping marks downstream circuits as well, we noted increased c-FOS in both the dorsal motor nucleus of the vagus (DMX) and rostral ventrolateral medulla (RVLM), major parasympathetic and sympathetic relay nuclei, respectively (Fig. 5C-D, S6C). This delineates a central pathway through which CYSLTR2 neurons access both parasympathetic and sympathetic outflow to the airways.

### Sympathetic neurons mediate the efferent arm of the vagal vascular leak reflex

We next asked whether the efferent arm of the reflex is sympathetic or parasympathetic. Pulmonary sympathetic fibers outnumber parasympathetic fibers near large pulmonary vessels (Fig S6D), positioning them as the likely effectors of CYSLTR2 neuron-mediated leak. We therefore optogenetically activated the vagosympathetic trunk in *Dbh-Cre;loxP-ChR2* and *Chat-Cre;loxP-ChR2* mice, which mark sympathetic and parasympathetic motor neurons respectively, and measured Evans blue extravasation (Fig. 5E). CHAT stimulation did not raise pulmonary dye beyond sham stimulation. By contrast, DBH stimulation disrupted the endothelial barrier to a degree comparable to CYSLTR2 activation, while sparing skin and abdominal organs innervated by more posterior sympathetic ganglia. Chemical sympathectomy blocked the Evans blue extravasation evoked by CYSLTR2 activation entirely (Fig. 5F). Because CYSLTR2 neurons also innervate other viscera, we asked whether this reflex is confined to the lung. Activation increased Evans blue extravasation in the heart and gut mucosa, while sparing the liver and skin (Fig. 5E-F). Direct stimulation of the vagosympathetic trunk engages cervical and upper thoracic outflow alone, whereas afferent activation recruits medullary relays with access to more posterior sympathetic ganglia, accounting for the difference in the organs affected. Vascular leak in diverse organs is therefore under direct sympathetic control, completing a reflex arc that runs from juxtacapillary afferents through medullary relays to sympathetic effectors.

## Discussion

Airway defense depends on the interplay of many vagal afferent classes, including dedicated pathways that respond to overinflation^42^, airway closure^4^, aspiration^3^, and various irritants^6,40,41^. Here, we describe a reflex initiated by an orphan population of vagal neurons that controls lung vascular leak. Recurrently activating these neurons causes fluid accumulation around the large vessels and bronchi, impairing lung mechanics independent of other airway insults. Single-cell profiling and functional silencing experiments confirm that these neurons are preferentially engaged during allergic inflammation and required for allergen-induced perivascular edema. These neurons act through brainstem relays that engage the sympathetic nerves of the upper chest, coordinating fluid balance across multiple organs, in contrast to the local mechanisms used by edema-promoting nociceptors in the skin. Together, we propose that CYSLTR2 neurons function as vascular state sensors, detecting mechanical and inflammatory compromise of the alveolar capillary bed and directing post-capillary extravasation, perhaps to support local repair and host defense. Under this model, the reflex is useful when engaged briefly and harmful when sustained, since the same extravasation that provisions the tissue also produces cuffing that eventually stiffens the lungs and disrupts breathing.

The subtype-resolved mapping of vagal afferents afforded by modern genetic tools is helping to clarify long-standing questions about lung C-fibers. For example, Paintal originally defined J receptors by their response to vascular mechanical stimuli, whereas later work defined them instead by their response to capsaicin. The two definitions are difficult to reconcile, because capsaicin-responsive neurons react only weakly to raised capillary pressure and lung inflation^65,66^. Furthermore, Paintal proposed that J receptor cause dyspnea (breathlessness) and exercise termination, yet human volunteers injected intravenously with capsaicin do not report dyspnea^73^. Our work may help explain these discrepant observations by showing that there are at least two subtypes of alveolar endings, that together account for features collectively ascribed to J receptors. Of the alveolar subtypes, N10 neurons (marked by *Vip*) respond to capsaicin have been shown to initiate airway hyperresponsiveness^45^, whereas N12 neurons (marked by *Cysltr2*) respond to mechanical stimuli but not TRP channel agonists and evoke leak from large pulmonary vessels. Other C fiber classes exist as well – for example, the airway wall mucosa is innervated by jugular nociceptors (marked by *Npy1r*, *Trpv1* and *Tac1* expression)^74^, which drive tachypnea and urge-to-cough^3,75^, and may also initiate vascular leak from the bronchial microvasculature. Natural insults likely engage more than one pathway at once, as we see for N10 and N12 neurons during OVA challenge, and the relative weights of pathway activation may account for different symptomatic outcomes seen across individuals or exposure types. It also leads to the tempting hypothesis that targeting one pathway could resolve some reflex outcomes while sparing others needed for eupnic breathing.

One implication of this work is that neural dysregulation could drive airway pathology rather than merely respond to it. Leakage of plasma from airway vessels is a recognized feature of many airway disorders, but has largely been viewed as a downstream consequence of inflammatory mediators acting directly on the endothelium^70,71^. Our results place both vagal and sympathetic neurons upstream of that process. Sympathetic control of pulmonary vascular leak is already recognized in the context of specific pathologies, like neurogenic and high-altitude pulmonary edema, but the evolutionary purpose of this adrenergic sensitivity was unclear. In those cases, sympathetic neurons are thought to evoke edema through changes in hydrostatic pressure, redistribution of blood volume, cardiac effects, reduced clearance, or direct effects on endothelial permeability^72^. In aggregate, we suggest that the sympathetic nervous system is not simply a source of pathological catecholamine excess, but an effector through which sensory information about the state of the lung can be translated into coordinated changes in organ fluid balance.

Our work also raises the possibility that neurotherapeutics could provide targeted benefit for conditions associated with excessive peribronchovascular leak. Cysteinyl leukotrienes themselves have well known roles in airway allergic disease, with CYSLTR1 antagonists in clinical use and CYSLTR2 variants associated with asthma and aspirin-exacerbated respiratory disease^73–75^. Our results suggest that these and other inflammatory mediators may also contribute to disease by engaging neural circuits that drive lung pathology, representing a new direction for clinical investigation. From these combined results, we conclude that CYSLTR2 vagal neurons constitute a neural axis controlling lung fluid balance in both health and disease.

## Acknowledgements

We thank Jayaraj Rajagopal, Stephen Liberles, Adam Haber, Michael Schappe, Chen Ran, Marito Hayashi, and members of the Prescott lab for manuscript comments. We thank Isaac M. Chiu for sharing the *Cysltr2-ires-Cre* mouse line. We thank Keith Murray for providing administrative support. Animal care and husbandry was provided by MIT Division of Comparative Medicine (DCM), histological processing and hematoxylin and eosin staining was provided by the Histology Core of the Koch Institute’s Robert A. Swanson (1969) Biotechnology Center, sequencing reagents and support were provided through the Biology of Adversity Project at the Broad Institute of MIT and Harvard, and additional computational support was provided the MIT BioMicro Center.

## Funding

This work was supported by a Child Health Research Award from the Charles H. Hood Foundation, by the Pew Biomedical Scholars Program in Biomedical Sciences, and by the Freedom Together Foundation. The work related to H&E staining was supported in part by the Koch Institute Support (core) Grant P30-CA14051 from the National Cancer Institute. EKR is a K. Lisa Yang Brain-Body Fellow, EI is a MIT HEALS Graduate Fellow, and BU is supported in part through the Biology of Adversity Project.

## Author contributions

EKR and SLP conceived the study, and EKR, SLP, and BU designed the experiments. EKR executed all mouse experiments and data collection. EKR, EI, BSO, and JL helped with ELISAs and vagal tracing. EKR, JM, and NB performed histology and imaging. EKR, BU and SLP collected samples for single cell RNA-sequencing, and EKR, BU, and SLP performed sequencing analysis. Formal analysis and development of MATLAB and Python scripts were performed by EKR, KS, BU, and SLP. Figures were generated by EKR, BU, SLP. The manuscript was written by EKR, BU, and SLP, and was reviewed and approved by all authors. Funding acquisition was provided by SLP.

## Competing interests

The authors declare that they have no competing interests.

## List of Supplementary Materials

Materials and methods

Table S1

Table S2

Table S3

Fig S1

Fig S2

Fig S3

Fig S3

Fig S4

Fig S5

Fig S6

Fig S7

Movie 1

Movie 2

Link to Github page: https://github.mit.edu/Prescott-Lab/bumans_OVA_manuscript

## Material & Methods

### Mice

All animal husbandry and procedures were performed in compliance with institutional animal care and use committee guidelines. Age-matched 8-11 week-old male littermate mice were used for all experiments unless otherwise noted. The following mouse transgenic lines were used: *Gabra1-ires-Cre* (gift from the Liberles lab, Harvard Medical School); *Npy1r-Cre* (Jackson Laboratory, 030544); *Kcng4-Cre* (Jackson Laboratory, 029414); *Vglut2-ires-Cre* (Jackson Laboratory, 029414); *Calb1-ires2-Cre* (Jackson Laboratory, 028532)*; Cysltr2-ires-Cre* (gift from the Isaac Chiu, Harvard Medical School); *Piezo2-eGFP-ires-Cre* (Jackson Laboratory, 027719), and *Vip-Cre* (Jackson Laboratory, 010908). Reporter/effector lines include *Ai14* (*loxP-tdTomato*; Jackson Laboratory, 007908), *Ai32* (*loxP-ChR2*; 012569), and *loxP-DTR* (Jackson Laboratory, 007900). Calcium imaging was performed on neurons from *Snap25-2A-GCaMP6s* mice (B6.Cg-Snap25tm3.1Hze/J; Jackson Laboratory, 025111), which express the genetically encoded calcium indicator GCaMP6s pan-neuronally from the endogenous *Snap25* locus, crossed to *Cysltr2-ires-Cre* and a *Cre*-dependent *Rosa26-loxP-tdTomato* reporter so that *Cysltr2*-lineage neurons could be identified by red fluorescence within the same imaging field. Wildtype controls were C57BL/6J (Jackson Laboratory, 000664).

### Ovalbumin (OVA) challenge

Male mice aged 8-11 weeks were sensitized by intraperitoneal (i.p.) injection of ovalbumin (OVA; 50μg/mouse; Sigma-Aldrich, A5503) with aluminum hydroxide adjuvant (1.6mg/mouse; Alhydrogel 2%, InvivoGen) on days 0 and 7. On days 16–18, mice were exposed to aerosolized 1% OVA (2 mL total volume) in 1 L/min oxygen flow for 30 minutes during the light cycle. Control animals received aluminum hydroxide adjuvant (1.6mg/mouse; Alhydrogel 2%, InvivoGen, i.p.) and saline on days 0 and 7, then exposed to aerosolized saline with 1 L/min oxygen flow for 30 minutes on days 16–18. Animals were anesthetized for lung function testing or euthanized for tissue harvest at day 20 unless otherwise specified.

### Methacholine challenge and lung function testing

Mice were anesthetized with tribromoethanol (375 mg/kg). Upon reaching a surgical plane of anesthesia, animals were tracheostomized, and mechanically ventilated (SAR-1000, CWE) at 150 breaths/min with a tidal volume of 7.5 mL/kg. Each ventilated breath was programmed with 120ms for inspiration, 80ms breath-hold maneuver, and 200ms for the expiration. The diaphragm was transected to eliminate spontaneous breathing. Positive end-expiratory pressure was set to 20mmH₂O and 100% oxygen was supplemented during ventilation. Tracheal pressure was measured by connecting a tracheal cannula port to a differential pressure transducer. Airflow was measured using a pneumotachograph (Harvard Apparatus, 73-0981) connected to a differential pressure transducer (Harvard Apparatus, 73-0064). Data from both transducers was amplified (BIOPAC, DA100C) and recorded continuously at 2 kHz using AcqKnowledge Software (BIOPAC).

For methacholine bronchoprovocation assays, mice were given methacholine every two minutes after a 1 minute baseline at a rate of 0.2mL/min (Doses: 25, 50, 100, 200, 400, 800, 1250 mg/kg; USP) via retro-orbital injection.

### Airway resistance and compliance measurements

Respiratory mechanics were computed breath by breath with custom MATLAB code (MathWorks) using the constant-flow, end-inspiratory-occlusion formulation established for mechanically ventilated subjects^76^. Briefly, breath boundaries were defined from the volume signal. Airflow was smoothed with a third-order Savitzky-Golay filter, corrected for slow baseline drift by subtraction of a running median, and integrated to yield volume. Inspiratory and expiratory volume extrema marked the ends of inspiration and expiration, and the mode of the kernel density estimate of peak prominences across the recording defined the average tidal volume used to set detection thresholds. Three pressure landmarks were then identified per breath. Peak inspiratory pressure (PIP) was taken as the maximum tracheal pressure between successive end-inspiratory volume peaks, refined to the local maximum within a 20-sample window. The end-inspiratory plateau pressure (Pplat) and the end-expiratory pressure (PEEP) were taken from the first and second inflections of the second derivative of the filtered pressure trace following PIP. Breaths whose PIP was an outlier relative to a 20-breath moving mean, and breaths whose PIP-to-plateau interval exceeded twice the median interval, were excluded.

Airway resistance (Raw) for each breath was calculated as the pressure drop across the resistive component divided by the inspiratory flow,

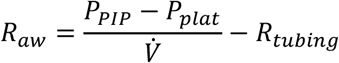

where *V̇* is the constant inspiratory flow in mL/min. The resistance of the cannula and ventilator tubing was measured in the absence of an animal (0.0101 mmH₂O per mL/min) and subtracted from every resistance value. Quasi-static compliance was calculated as the delivered tidal volume divided by the elastic pressure drop,

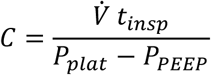

where *t_insp_* is the fixed inspiratory flow time (taken as the interval from end-expiration to PIP), so that the product *V̇t_insp_* is the volume delivered during inspiratory flow. Resistance is reported in mmH₂O/(mL/min) and compliance in mL/mmH₂O. For each methacholine dose, the 60-s window beginning at the time of delivery was analyzed, and the area under the resistance curve and the median compliance within that window are reported.

To quantify breath-to-breath instability of airway resistance, we performed a sliding-window spectral analysis of the detrended breath-by-breath resistance signal. A robust locally weighted trend (rlowess, span 5 breaths) was subtracted from this series to yield resistance residuals, isolating breath-to-breath fluctuations from the slow underlying trend. The residual series was then transformed into the time–frequency domain by a short-time Fourier transform (spectrogram; 30-breath Hamming window, 29-breath overlap, evaluated at the breath-series sampling rate), and the power spectrum was taken as the squared magnitude of the transform. At each time bin, a spectral root-mean-square (RMS) amplitude was calculated as the square root of the mean spectral power across all frequency bins, giving a breath-indexed trace of the RMS amplitude of the resistance residuals. For each methacholine dose, a 60-s analysis window beginning at that dose’s marker time was defined, and “RMS (AUC)” was computed as the trapezoidal integral of the spectral RMS trace over this window (time in seconds), yielding one cumulative RMS value per dose per animal that was used as the summary response in subsequent dose–response analyses.

Dose-response curves were compared between groups with a linear mixed-effects model with mouse as a random intercept, fit by maximum likelihood. An omnibus *P* value was obtained by likelihood-ratio comparison of Response ∼ Group × Dose + (1|Mouse) against Response ∼ Dose + (1|Mouse), with dose treated as a categorical factor. This test detects a difference in overall level, in curve shape, or both. Curves show group mean ± SEM (n mice per group given in each figure).

### Tissue collection and immunofluorescence

Mice were euthanized with pentobarbital and then cardiac perfused with ice-cold 4% paraformaldehyde (PFA) in saline. Lungs were inflated with 1mL of cold 4% PFA before dissection through the trachea at a rate of 1mL/min. Tissues were post-fixed overnight in 4% PFA at 4°C. Brains and lungs were cryoprotected in 30% sucrose, incubated in O.C.T. for 1 hour before freezing and sectioning.

For lung tracing experiments, thick (300-µm) cryosections were stained with primary and secondary antibodies as listed in Table S1 for 16-36 hours at 4°C. Samples were cleared using EasyIndex (LifeCanvas Technologies) and imaged on a Leica Stellaris confocal microscope.

Brain sections (40µm) were stained with primary and secondary antibodies as listed in Table S1.

Ganglia were processed using the EZ Clear protocol^77^ and stained with primary antibodies as listed in Table S1.

### Serum IgE quantification by ELISA

Total and OVA-specific IgE were measured in serum collected at day 20 of the challenge protocol. Mice were anesthetized with avertin (tribromoethanol; 375 mg/kg i.p.), the descending aorta was transected, and approximately 1 mL of blood was collected into serum separator tubes. Samples were held on ice until collection was complete for all animals in the cohort, then allowed to clot for 45 minutes before being centrifuged at 2,000 × g for 15 min at 4 °C. Serum was transferred to clean tubes and stored at −80 °C until assay, with no more than one freeze– thaw cycle.

Total IgE was measured with Mouse Total IgE ELISA Kit (RayBio Tech, ELM-IgE-1) and OVA-specific IgE with LEGEND MAX™ Mouse OVA Specific IgE ELISA Kit (Biolegend, 439807) following the manufacturer’s instructions. Samples were assayed in duplicate at [1:100] (total IgE) and [1:50] (OVA-specific IgE); dilutions chosen so that readings fell within the linear portion of the standard curve. Standards were run in duplicate across 7 serial dilutions on every plate. Absorbance was read at 450 nm on a ThermoFisher Varioskan Flash plate reader, and concentrations were interpolated from a four-parameter logistic fit. The limit of detection was 20.7 pg/mL (for Anti-OVA IgE ELISA) and 0.14ng/mL(for Total IgE ELISA); samples reading below it were assigned the limit of detection. Where samples were distributed across more than one plate, a pooled reference serum was included on every plate and used for normalization.

### Nodose ganglion injections and vagotomy

Animals were anesthetized using isoflurane (1-3%, delivered in oxygen) and placed in a supine position. Vagal ganglion injections were performed as previously described^3,5^. AAVs were mixed with Fast Green dye (10:1 ratio, to 0.1% final dye concentration) and injected into vagal ganglia in a volume of 140nl, using a Nanoject II injector (Drummond), and success determined visually by Fast Green Dye filling of the ganglia. Tracing experiments used a 1:1 titer ratio of *pAAV-Flex-tdTomato*(Addgene 28306-PHP.S, v247992, 1.9 × 10^13 gc/mL) and *pENN.AAV.CB7.CI.eGFP.WPRE.rBG* (Addgene 105542-AAV9, v36906, 2.3×10^13 gc/mL). Chemogenetic experiments used *AAV-hSyn-DIO-hM3D(Gq)-mCherry* (Addgene 44361-AAV9, v227973, 1.5×10^13 gc/mL) or *AAV-hSyn-DIO-hM3D(Gi)-mCherry* (Addgene 44362-AAV9, v241463, 1.9×10^13 gc/mL) viruses or *pAAV-Flex-tdTomato* virus for control experiments.

For vagotomy experiments, the vagus nerve was exposed following the same procedure as used for nodose ganglion injections. The vagal trunk was then resected from surrounding tissue, transected approximately 1mm below the SLN branchpoint and cauterized following unilateral nodose injection.

### Chemogenetic vagal manipulation

Following ganglion injection of *AAV-hSyn-DIO-hM3D(Gq)-mCherry* or *AAV-hSyn-DIO-hM3D(Gi)-mCherry* viruses, mice recovered for 1 week before initiating OVA sensitization. Compound 21 (C21; 3 mg/kg) was administered via intraperitoneal injection twice daily (8-hour interval) on days 16–18, on concurrent days with the OVA aerosol challenge. Methacholine measurements were performed on day 20. Additionally, D18 (2 hours after last C21 dose) and D27 timepoints were used for lung structure analyses.

### Brainstem tracing and c-Fos mapping

For brainstem tracing, male *Cysltr2-ires-Cre* mice received unilaterally injections of a viral tracing cocktail containing *pAAV-Flex-tdTomato* and *pENN.AAV.CB7.CI.eGFP.WPRE.rBG* (1:1) into the nodose ganglion, as described above. Mice were allowed to recover for 4 weeks before pentobarbital euthanasia, followed by tissue collection and sectioning as described above.

For c-Fos mapping experiments, male *Cysltr2-ires-Cre* mice (*Cysltr2*-Gq) and C*re-*negative littermate controls received injections of *AAV-hSyn-DIO-hM3D(Gq)-mCherry* into the nodose ganglion. Mice were allowed to recover for 4 weeks before intraperitoneal administration of C21 (3 mg/kg). Ninety minutes after C21 administration, mice were euthanized with pentobarbital overdose, and tissues were collected as described above. Brainstem sections were co-stained with antibodies against c-Fos and mCherry, provided in Table S1.

### Single-cell RNA sequencing

Nodose/jugular ganglia were harvested from 80 adult male C57BL/6J mice in two batches, corresponding to days 20 and 95 after initial ovalbumin exposure. In each batch, four replicate samples of ovalbumin- or PBS-treated animals, each consisting of ganglia pooled from five animals, were collected. Each pooled sample was dissociated in 1 mL of 1:1 Dulbecco’s Modified Eagle’s Medium/Ham’s F12 media (DMEM; Thermo Fisher Scientific, 11039047) supplemented with Liberase TH (0.22 mg/mL) and DNase (100 U/mL) at 37°C for 30 min.

Ganglia were triturated with a P200 tip, filtered through a 70 µm cell strainer and pelleted (300 × g, 10 min, 4°C). Cells were fixed overnight according to GEM-X Flex protocols (10x Genomics, CG000782) and stored at −80°C. Fixed cells were processed for sequencing in a single multiplexed batch using the 10X Genomics Flex v2 platform, according to manufacturer instructions. Libraries were sequenced on an Illumina NovaSeq X platform.

Fastq files were aligned and samples were demultiplexed using cellranger software. After processing with CellBender, feature barcode matrices were preprocessed and filtered to exclude cells with fewer than 200 UMIs. Coarse clustering (resolution 0.3) was used to isolate sensory neuron populations and detect doublets using scDblFinder^78^, and nodose and jugular sensory neurons, identified by cluster-wide expression of *Phox2b* or *Prdm12*, were independently subclustered (resolutions 0.35 and 0.2, respectively) using Seurat (v5)^79^ prior to further analysis. Data were compared to prior vagal single-cell datasets using Seurat’s MapQuery function. Treatment-responsive cell populations were identified using scDist^80^, modeling ovalbumin vs. PBS treatment status as a fixed effect and replicate sample as a random effect. Benjamini-Hochberg false discovery rate estimates were used to identify significantly altered subtypes (N10 and N12 only). Separately, we identified differentially-expressed genes in each cell cluster independently, using DeSeq2^81^ to model pseudobulk expression (calculated for each sample/cluster combination) as a function of treatment condition. In both scDist and DeSeq2 analyses, we compared ovalbumin- and PBS-treated samples at the early and late time points independently to avoid potential confounding influences of age or sample collection batch. The fgsea package^82^ was used to test for enrichment of gene sets obtained from MSigDB (Hallmark, GO-Biological Processes, Reactome). Gene sets with fewer than 15 or more than 500 genes expressed in a subtype were excluded, and the Wald statistic obtained from differential expression testing was used to rank genes for enrichment testing.

### Optogenetics

Mice were anesthetized with urethane (2 mg/g) for spontaneous breathing experiments or tribromoethanol (375 mg/kg) for ventilated conditions. The nodose/jugular ganglion was exposed, and an optic fiber (400 µm core; Thorlabs) was positioned over the vagal trunk. Light stimulation (10–50 Hz, 5-ms pulses, 10–30 s duration) was delivered via a DPSS laser system (473 nm, 150 mW, Ultralaser). Heart rhythm was measured via EKG, which was recorded with two needle electrodes placed subcutaneously on the right forepaw and left hindpaw and amplified with a differential amplifier (A-M Systems), with a baseline needle in the right hindpaw to reduce noise. For spontaneous breathing experiments, tracheal pressure was measured using an amplifier-coupled pressure transducer (Biopac) cannulated into the trachea and acquired at 1 kHz (MP150, Biopac). Pressure was detrended (Gaussian moving average, 5,000 samples), integrated over time by cumulative trapezoidal integration to obtain relative lung volume, and the integrated signal was detrended again (Savitzky–Golay, 10,000 samples). The modal tidal volume of each recording was estimated from a kernel density estimate of breath amplitudes (peak prominences), and a breath was scored if the lung-volume excursion reached ≥30% of this modal tidal volume with a minimum inter-breath interval of 0.2 s; end-inspiratory maxima and end-expiratory minima defined each breath. Respiratory rate was calculated from consecutive end-inspiratory intervals, tidal volume was taken as the end-inspiratory peak prominence of the integrated trace, heart rate was calculated from R-wave intervals of a baseline-corrected, rectified EKG, and all variables were expressed as the percent change between the 20 s preceding and the 10 s following each stimulus. For ventilated experiments, breathing was recorded and airway resistance and dynamic compliance were calculated from tracheal pressure and flow exactly as described above (see Airway resistance and compliance measurements), and, as for spontaneous breathing, responses were expressed as the percent change between the 20 s preceding and the 10 s following each stimulus.

### Evans Blue vascular permeability assay and Wet-to-Dry Ratio Measurements

Mice were anesthetized with tribromoethanol (375 mg kg⁻¹, i.p.) 5 min before retro-orbital injection of Evans Blue (50 mg kg⁻¹ in saline). Evans Blue was allowed to circulate for 10 min before optogenetic stimulation (10Hz for motor activation and 50Hz for sensory activation, 5-ms pulses, 10 s on/20 s off) for 10 min. Mice were then cardiac perfused with cold saline containing 2 mM EDTA for 10 min. Lung, upper gut (mm length), left liver lobe, skin (mm² area), and heart samples were homogenized in 500 μl formamide and incubated at 60°C for 12–16 h. Samples were centrifuged, and Evans Blue absorbance in the supernatant was measured at 620 and 700 nm. Absorbance at 700 nm was subtracted from the 620-nm measurement to correct for heme background. Corrected absorbance values were normalized to the mean of control samples within each experimental cohort.

For wet-to-dry weight measurements, mice were anesthetized with tribromoethanol (375 mg kg⁻¹, i.p.) 15 min before they underwent the same optogenetic stimulation conditions as described above. Mice were then cardiac perfused with cold saline containing 2mM EDTA for 10 mins. The postcaval lung lobe, right atrium, gut (mm length), skin (mm² area), and mid liver lobe were collected and then weighed immediately. Samples were dried at 60°C for 72 h before dry weight was determined.

### H&E Staining

Lung tissues were fixed, processed, and sectioned at 10-25 μm thickness. Sections were stained with hematoxylin and eosin (H&E) using standard histological methods by the MIT Koch Histology Facility. Whole-slide images were acquired using a digital slide scanner for histological assessment and image analysis.

### Stereology, cuff measurements and nuclei density measurements

#### Image acquisition and region definition

H&E-stained 10-25 µm-thick lung lobe sections were imaged as whole slides and annotated in QuPath v0.7.0. For each section a reference annotation delimiting the lobe was drawn, together with exclusion annotations covering folds and section-edge artefact; the effective reference is the annotation minus exclusions. Regions were exported at 2 µm/px as co-registered RGB, reference-mask and exclusion-mask TIFFs. Sections were excluded from futher analysis if the effective reference was < 2 mm², if it was < 10% of the median region area for its slide, if < 5 sampling frames were placed, or if V_v_(area) > 0.85 (indicating fold, clot or debris).

#### Stain deconvolution and tissue segmentation

RGB images were separated into haematoxylin (H) and eosin (E) optical density by Ruifrok–Johnston color deconvolution^83^ using the “H&E 2” vectors (H = 0.490, 0.769, 0.410; E = 0.046, 0.842, 0.537), with OD = −log₁₀((I + 0.5)/255.5). Within the effective reference, pixels with OD(H) + OD(E) ≥ 0.15 were classified as tissue and the remainder as airspace. This single threshold defines every quantity below.

#### Nuclei counts

Nuclei were segmented in QuPath using InstanSeg extension v0.1.1 at the native 0.504 µm/px resolution, on a systematic-uniform-random sample of 300 µm frames using the same lattice and seed as the stereological sampling below. InstanSeg counts are reported per mm² of parenchyma (effective reference minus large lumens).

#### Stereology

Stereology metrics were calculated using custom Python scripts. Airspace components ≥ 200 µm equivalent diameter (large lumens) and dense tissue folds (100 µm tiles with mean OD ≥ 0.35 and ≥ 60% of pixels above that value) were excluded from the parenchymal compartment. Sampling used 300 µm frames on a 600 µm lattice with a seeded random offset, a 25 µm point grid, and six randomly rotated 250 µm line probes per frame. We report tissue volume fraction V_v_ by point counting, air–tissue surface density S_v_ = 2·I/L, interstitial thickness τ = 2·V_v_/S_v_, and mean linear intercept MLI = 4·(1 − V_v_)/S_v_. Stain density is reported as mean OD per unit effective reference area. Local tissue density was additionally measured as the distribution of tissue fraction in 200 µm windows lying wholly inside the reference, with lumens and folds treated as excluded rather than empty pixels.

#### Airway/blood vessel detection and classification

To identify airways and blood vessels, candidate airspace components were first proposed permissively (60–4,000 µm equivalent diameter, inscribed radius ≥ 20 µm, wall occupancy ≥ 0.55 within 20 µm and ≥ 0.25 at 20–40 µm, evaluated only where the wall shell lies inside the reference). Candidates were then classified as ‘vessel’, ‘airway’, or ‘neither’ using a random forest (600 trees, balanced class weights). Twelve geometric and stain features were used, including rim OD expressed as a ratio to each section’s own median tissue OD so that staining batch cannot influence selection. The model was trained on 1172 blind-annotated objects from 187 sections and assessed by 5-fold cross-validation grouped by section, giving an overall accuracy of 0.84 (precision and recall 0.62 and 0.66 for airways, 0.66 and 0.55 for vessels). Lumens were analysed at P(vessel) + P(airway) ≥ 0.60; vessels were defined as P(vessel) > 0.5; airways were defined permissively as P(airway) > P(vessel). Detection was verified against a per-region overlay generated from the same masks used for measurement. The trained classifier, its SHA-256 digest and the blinded label set are archived with the code.

#### Cuff thickness calculations

Collar thickness was measured as the Euclidean distance from each point on the one-pixel shell outside a detected lumen to the nearest airspace that is not itself a detected lumen, sampled at full resolution and suppressed within 10 µm of the reference boundary. Per lumen we report the median of this distribution. Analyses are stratified by lumen caliber using both area-equivalent and maximum-inscribed diameter, since the former overstates the bore of elongated and branched profiles. For caliber sweeps, a single omnibus test per lumen class was obtained by exhaustive enumeration of all 9,240 within-block reassignments of the treatment label, giving an exact P value, with a family-wise-corrected max|t| statistic identifying the caliber band contributing most. Negative results are reported as effect sizes with 95% confidence intervals rather than as P > 0.05.

#### Statistics

N refers to animals throughout, and no test treats sections, lumens or pixels as independent. Per-section values were aggregated to the animal as region-area–weighted means; collar thickness was aggregated as the median of per-lumen medians. Group comparisons used two-sided Welch t-tests on per-animal values, and every contrast is additionally reported from an OLS model including tissue volume fraction as a covariate, to account for differences in inflation states. Where both arms of a contrast were present in more than one staining round or slide block, that factor was included as a fixed effect and the blocked estimate is the one reported; contrasts in which a group is absent from some strata are flagged as confounded rather than pooled. Timepoints were never concatenated.

#### Software versions

Analysis used Python 3.10.12 with NumPy 2.2.6, pandas 2.3.3, SciPy 1.15.3, scikit-image 0.25.2, scikit-learn 1.7.2, statsmodels 0.15.0 and Matplotlib 3.10.9;

### UEA-1+ area/ airspace measurements

UEA-1 staining was quantified using custom MATLAB scripts. Images were analyzed in a fixed, randomly permuted order (fixed seed, saved to disk) to minimize bias. For each field, the full airway (lumen, wall, and epithelium) was manually outlined on the structural reference channel to generate a binary airway mask; masks were saved and all measurements were computed from the saved masks and raw images. Mucus-positive pixels were defined by a fixed intensity threshold (>0.10 of the normalized intensity range) applied to the UEA-1 channel, and mucus density was calculated as UEA-1–positive pixels within the airway mask per 1,000 airway pixels. A 300-dpi quality-control overlay of the airway boundary and UEA-1 signal was generated and reviewed for every image.

### CD45 area/ DAPI area measurements

CD45⁺ immune infiltration was quantified in whole lung lobe sections (5× confocal, channel 1 = DAPI, channel 2 = CD45) using a custom MATLAB script. Images were analyzed blinded, with coded identifiers assigned in randomized order and the identifier-to-filename key withheld until all images were complete. Folds and artifactual regions were manually outlined and excluded, and lung tissue was defined as DAPI⁺ pixels (intensity > 0.015) outside excluded regions.

CD45⁺ pixels were identified with a local adaptive threshold equal to the Gaussian-smoothed local background (σ = 10 px) plus 0.5 × the local Gaussian-smoothed absolute deviation from that background, subject to a hard minimum of 0.05 normalized intensity; objects < 3 px were discarded. CD45⁺ burden was expressed as CD45⁺ pixel area divided by DAPI⁺ lung area, with CD45⁺ object count per lung area as a secondary measure. Every image was reviewed against a six-panel segmentation QC montage, and any threshold adjustment was made blinded and logged.

### Color deconvolution of Evans blue and blood signal

RGB images of whole lung lobes were separated into component channels in Fiji using the Colour Deconvolution plugin, which implements the stain-unmixing method of Ruifrok and Johnston (ref). Because Evans blue and hemoglobin are not standard histological stains, a custom three-component stain vector matrix was defined empirically: regions containing predominantly Evans blue, predominantly blood, or unstained background tissue were sampled from control images, and their mean RGB values were entered as the three stain vectors (exact R, G, B coordinates for each component are given in table S2). The identical vector matrix was applied to every image across all experimental groups. Deconvolution produced separate 8-bit Evans blue and blood channels, with the third component capturing residual background. All lobes were imaged under identical illumination, exposure, and white-balance settings.

### *In situ* Hybridization

Fluorescence in situ hybridization was performed using the HCR v3.0 protocol (Molecular Instruments) to detect *Cysltr2* and *ChR2* transcripts. Tissue sections (14um) were fixed, permeabilized, and incubated with target-specific HCR probe sets against *Cysltr2* and *ChR2*, followed by hybridization with fluorescently labeled HCR hairpins according to the manufacturer’s protocol. Sections were washed and mounted for fluorescence imaging. Probe sets were obtained from Molecular Instruments.

### Calcium Imaging of dissociated vagal sensory neurons

Nodose/jugular ganglia were resected into ice-cold DMEM, cleaned of connective tissue and nerve rootlets, and pooled on ice; ganglia from 1 mouse yielded sufficient neurons for several rounds of imaging in a single day. Ganglia were settled by gravity, the DMEM was aspirated, and ganglia were digested in DMEM containing 0.11 mg/mL Liberase TH (collagenase I/II– thermolysin blend; Roche 5401135001) and 0.004% DNase I (Worthington LS002007 or recombinant DNase I, Roche 04716728001) for 45 min at 37 °C with rotation, then pelleted (300 × g, 5 min, 4 °C, swinging-bucket rotor).

Ganglia were resuspended in ice-cold 1× low-ovomucoid inhibitor solution (0.15% BSA [Sigma-Aldrich A8806], 0.15% ovomucoid trypsin inhibitor [Worthington LS003086], and 0.004% DNase I in DMEM, pH 7.4, 0.22-µm filtered) and mechanically dissociated by ∼15–20 gentle passages with a P200 pipette held flat against the tube bottom, avoiding bubbles, until the medium became cloudy. The suspension was diluted in the same solution, filtered through a pre-wetted 70-µm strainer, pelleted (300 × g, 5 min, 4 °C), washed once more in low-ovomucoid solution, and re-pelleted.

Neurons were resuspended at high density in low-ovomucoid solution and spot-seeded as small droplets onto the center of dry, pre-coated poly-D-lysine/laminin 12-mm cover glasses (Neuvitro GG-12-laminin) to concentrate cells within a single field of view. Cells attached for 10 min at 37 °C in a humidified incubator, after which pre-warmed medium was layered on top to prevent desiccation.

For imaging, cover glasses were mounted in a perfusion chamber and superfused continuously with room temperature HBSS containing Ca²⁺ and Mg²⁺, which is required for robust agonist-evoked responses. Solutions were delivered by gravity flow at a constant ∼7 mL/min with gravity evacuation, with flow rate consistent across all solutions. Epifluorescent images were acquired at 8 Hz (125 ms per frame) through a 5× air objective, which captured the entire seeded field in a single frame; tdTomato was imaged in a separate channel to assign *Cysltr2*-lineage identity to each region of interest. All stimuli were prepared in HBSS with Ca²⁺ and Mg²⁺ and applied sequentially by switching the perfusion line, with HBSS washout between applications: histamine (Millipore Sigma, 1 µM), LTC_4_ and LTD_4_ cocktail (cysteinyl leukotrienes; Cayman Chemical Company, 100nM), allyl isothiocyanate (AITC; Millipore Sigma, 100 µM), bradykinin (Millipore Sigma, 1 nM), Yoda1 (Tocris, 10 µM), serotonin (Millipore Sigma, 10 µM), and capsaicin (Millipore Sigma, 2 µM). KCl (Sigma Aldrich, 50 mM) was applied last in every experiment as a positive control to identify all viable neurons.

### Sympathetic ablation

Mice received 6-hydroxydopamine (6-OHDA; Cayman Chemical Company, 100 mg kg⁻¹ on day −6 and 250 mg kg⁻¹ on day −4) in 0.1% ascorbic acid (Sigma Aldrich) in saline via intraperitoneal injection. Evans Blue vascular permeability was assessed on day 0 as described above.

## Supplementary Figure Legends

**Fig. S1.**
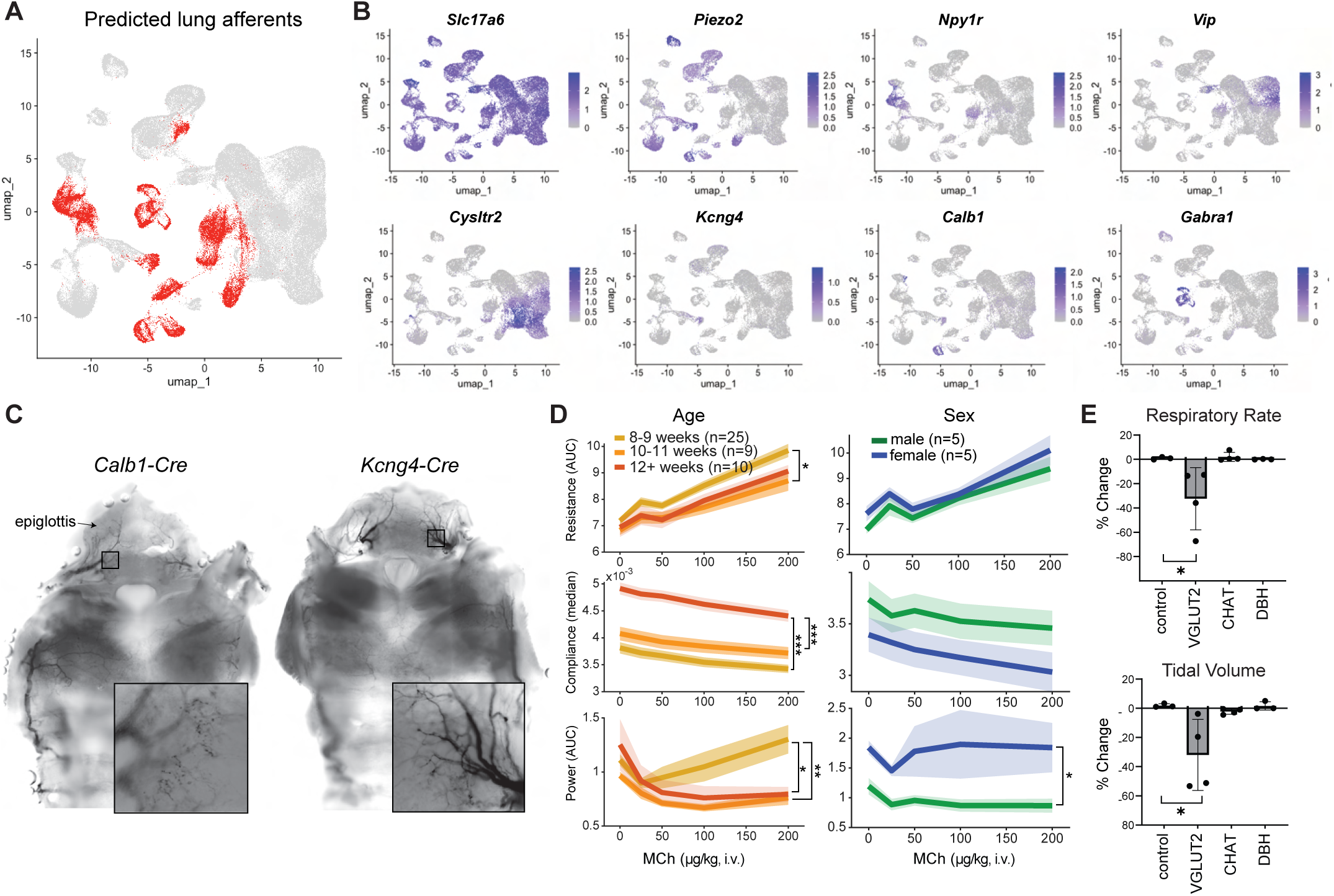
Genetic access to lung-innervating vagal sensory populations, and validation of airway mechanics across age and sex. **(A)** UMAP showing predicted lung afferents based at least one of three published datasets^3,13,16^. **(B)** UMAPs showing marker gene expression across vagal sensory neuron subsets. **(C)** Top: larynx whole-mount preparations comparing *Calb1-ires-Cre* and *Kcng4-Cre* tracing. Bottom: higher magnification of laryngeal terminals. **(D)** Resistance (AUC), compliance (median), and wave power during MCh challenge (i.v.; 25, 50, 100, 200 μg/kg). Left: by age, 8 to 9 weeks (yellow, n = 25), 10 to 11 weeks (orange, n = 9), and ≥12 weeks (red, n = 10). Right: by sex, male (green, n = 5) and female (blue, n = 5). Omnibus model with dose as a categorical factor, group × dose interaction reported. **(E)** Change in respiratory rate and tidal volume during vagal optogenetic stimulation in freely breathing mice expressing *ChR2* from the indicated *Cre* drivers. n = 3–4 mice per line; two-tailed unpaired t test; *P < 0.05, **P < 0.01, ***P < 0.001, ****P < 0.0001. Data are means ± SEM.

**Fig. S2.**
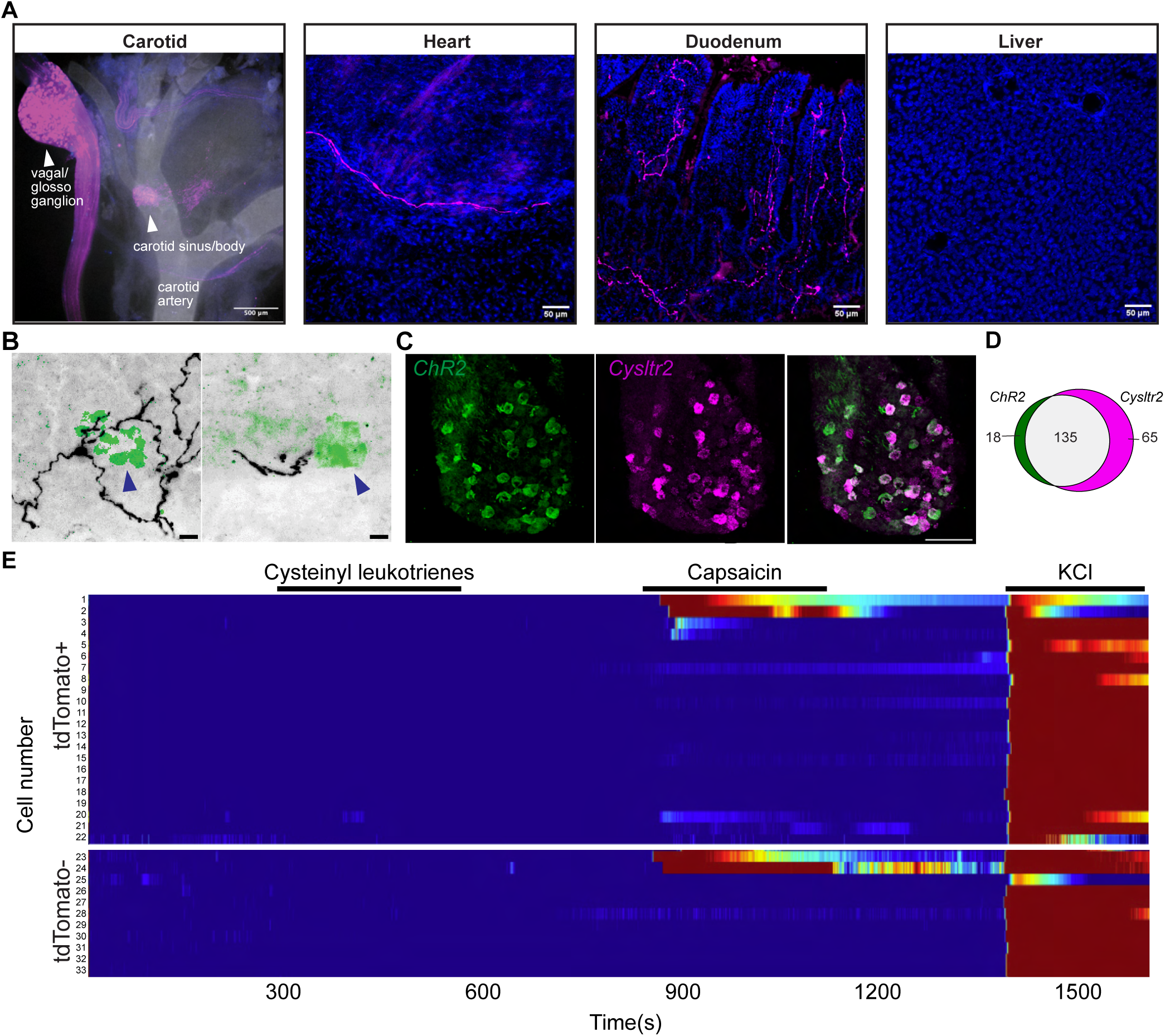
Cysltr2⁺ vagal neuron projection sites outside the lungs. **(A)** Confocal images of *Cysltr2*⁺ vagal fibers in the indicated organs (300-μm cryosections, anti-tdTomato immunostaining) from *Cysltr2-ires-Cre* mice injected in the nodose ganglia with *AAV-flex-tdTomato*. Scale bars are indicated for each image. **(B)** *Cysltr2*+ fibers (black) innervate neuroepithelial bodies (green) marked by blue arrows. Scale bars, 10 μm. **(C)** Representative image showing FISH labeling on a 14 μm section of nodose for *ChR2* (green) and *Cysltr2* (magenta) probes manufactured by Molecular Instruments. Scale bar, 100 μm. **(D)** Venn diagram shows the number of cells expressing either probe alone or both probes, quantifying the overlap in expression at the single-cell level. **(E)** Heatmap of ΔF/F₀ over time for tdTomato⁺ (22 out of 22 cells) and tdTomato-neurons (11 out of 39) during sequential perfusion of LTC_4_ and LTD_4_ cysteinyl leukotriene cocktail, capsaicin, and KCl and ranked by capsaicin response.

**Fig. S3.**
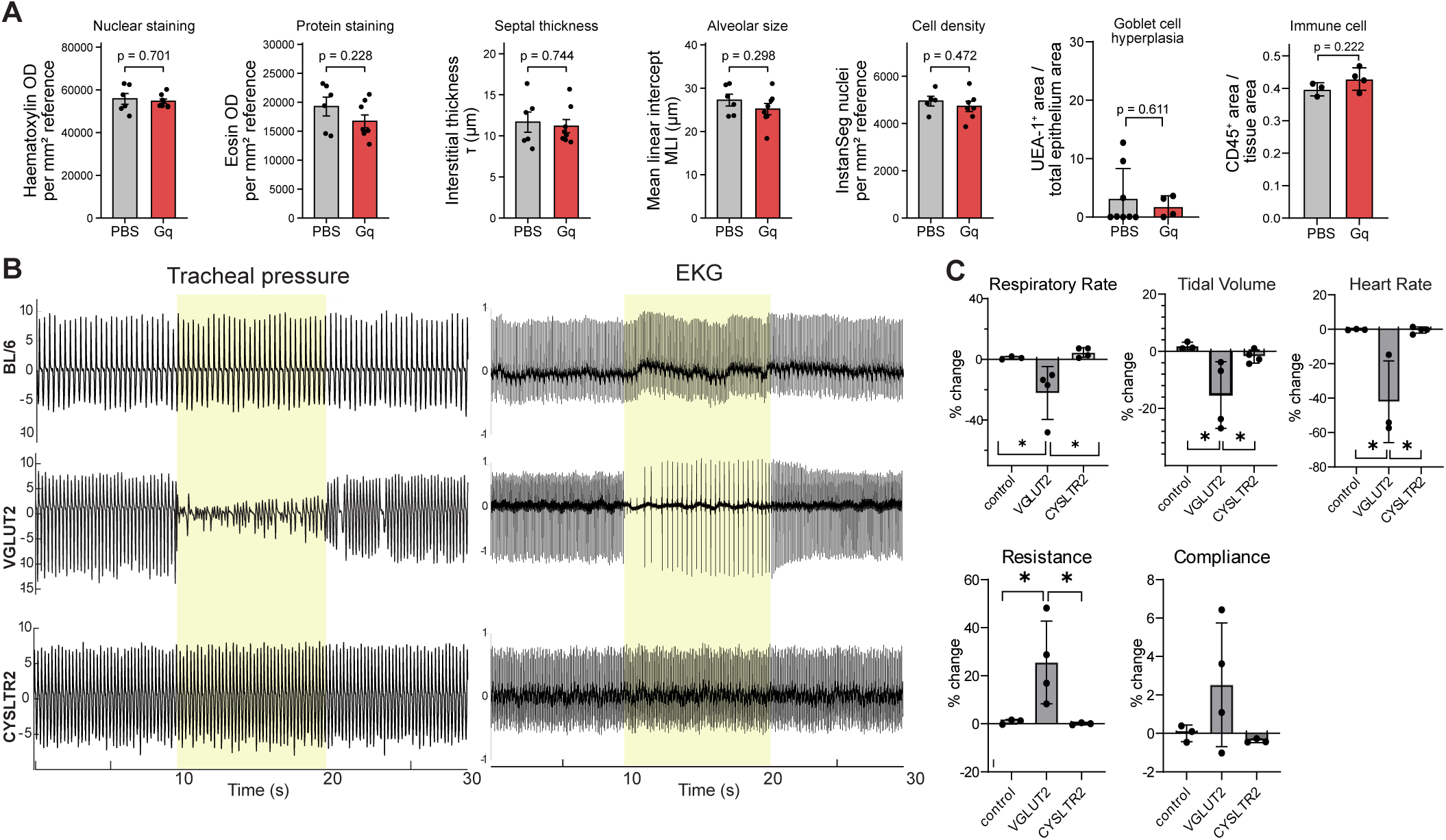
*Cysltr2*⁺ vagal neuron activation selectively affects pulmonary vascular physiology. **(A)** Nuclear staining, protein staining, septal thickness, alveolar size, cell density, goblet cell hyperplasia, and immune cell infiltration in *Cysltr2-ires-Cre* mice injected in the NJP ganglia with *AAV-Flex-tdTomato* (control) or *AAV-hSyn-DIO-hM3D(Gq)-mCherry* (Cysltr2-Gq). n = 3–8 mice per group; Welch’s two-tailed unpaired t test. **(B)** Representative tracheal pressure (left) and heart rate EMG (right) traces during optogenetic stimulation (yellow overlay) in *Cre*-negative, *Vglut2-Cre;loxP-ChR2*, and *Cysltr2-Cre;loxP-ChR2* mice. **(C)** Respiratory rate, tidal volume, resistance, compliance, and heart rate for the indicated *Cre* lines. n = 3–4 mice per line; two-tailed unpaired t test; asterisks as in Fig. S1E. Data are means ± SEM.

**Fig. S4.**
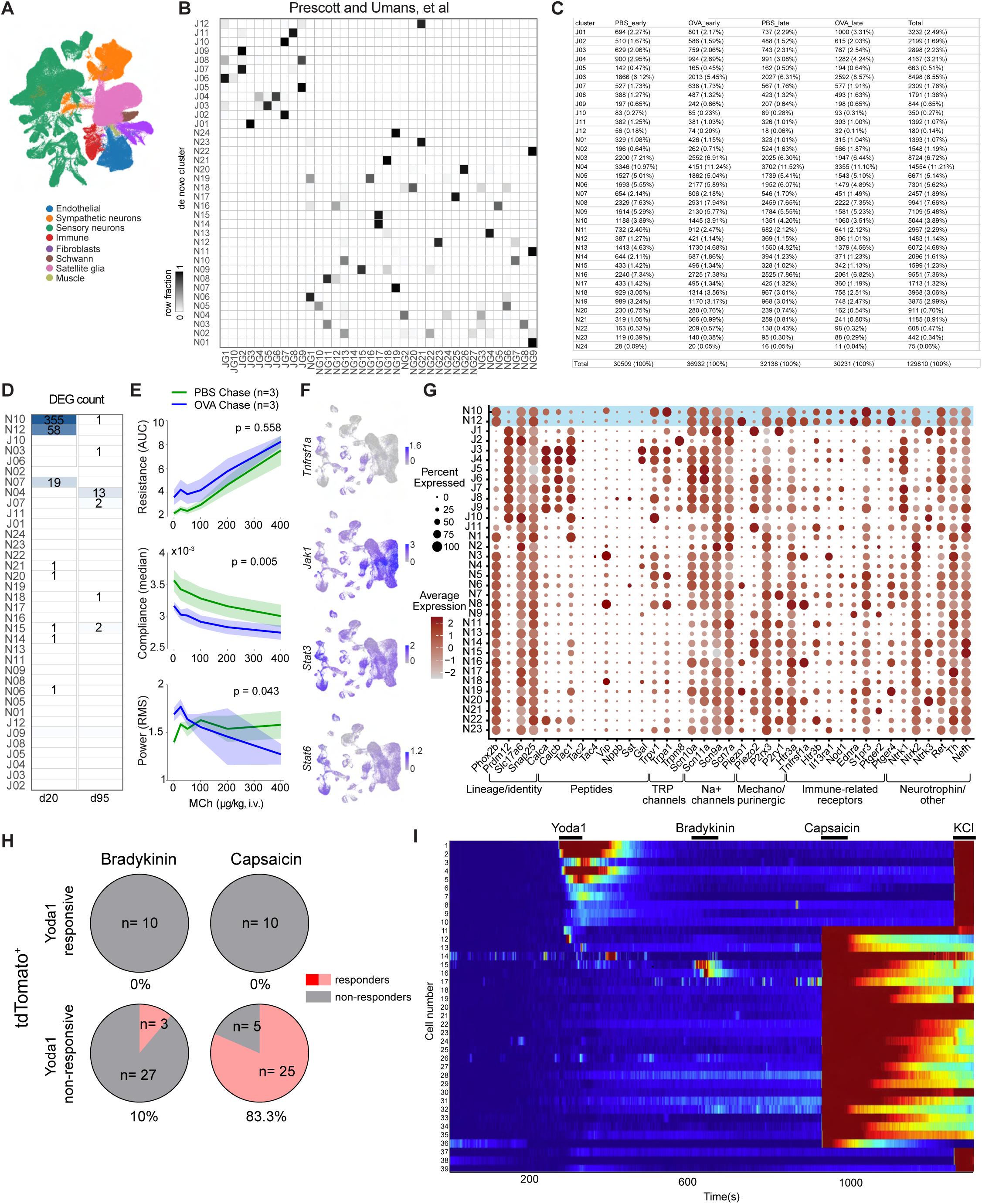
Extended single-cell transcriptomic analysis of vagal sensory neurons during allergic inflammation. **(A)** UMAP showing cell type identity for all cells collected for scRNA-seq. **(B)** Cluster correspondence matrix comparing de novo clusters (y axis) with Prescott et al. clusters (x axis); black indicates transcriptionally matched pairs. Each de novo subtype mapped to a single published cluster. **(C)** Cell counts per de novo cluster, split by condition (PBS early, OVA early, PBS late, OVA late) and total, with the percentage of each group’s total indicated. **(D)** DEG counts per cluster at day 20 and day 95. **(E)** Resistance (AUC), compliance (median), and wave power during MCh challenge (i.v.; 25, 50, 100, 200, 400 μg/kg) in PBS chase (green, n = 3) and OVA chase (blue, n = 3) mice. P-values calculated with omnibus model as in Fig. S1D. **(F)** UMAPs showing expression of the indicated genes. **(G)** Dot plot showing the percentage of cells expressing each gene and mean expression level across vagal sensory subtypes. **(H)** Proportion of tdTomato⁺ neurons responding to Yoda1 and, among those, the proportion responding to bradykinin or capsaicin; n and percentages are shown on each chart. **(I)** Heat map of ΔF/F₀ over time for tdTomato⁺ neurons during sequential perfusion of Yoda1, bradykinin, capsaicin, and KCl. Data are means ± SEM.

**Fig. S5.**
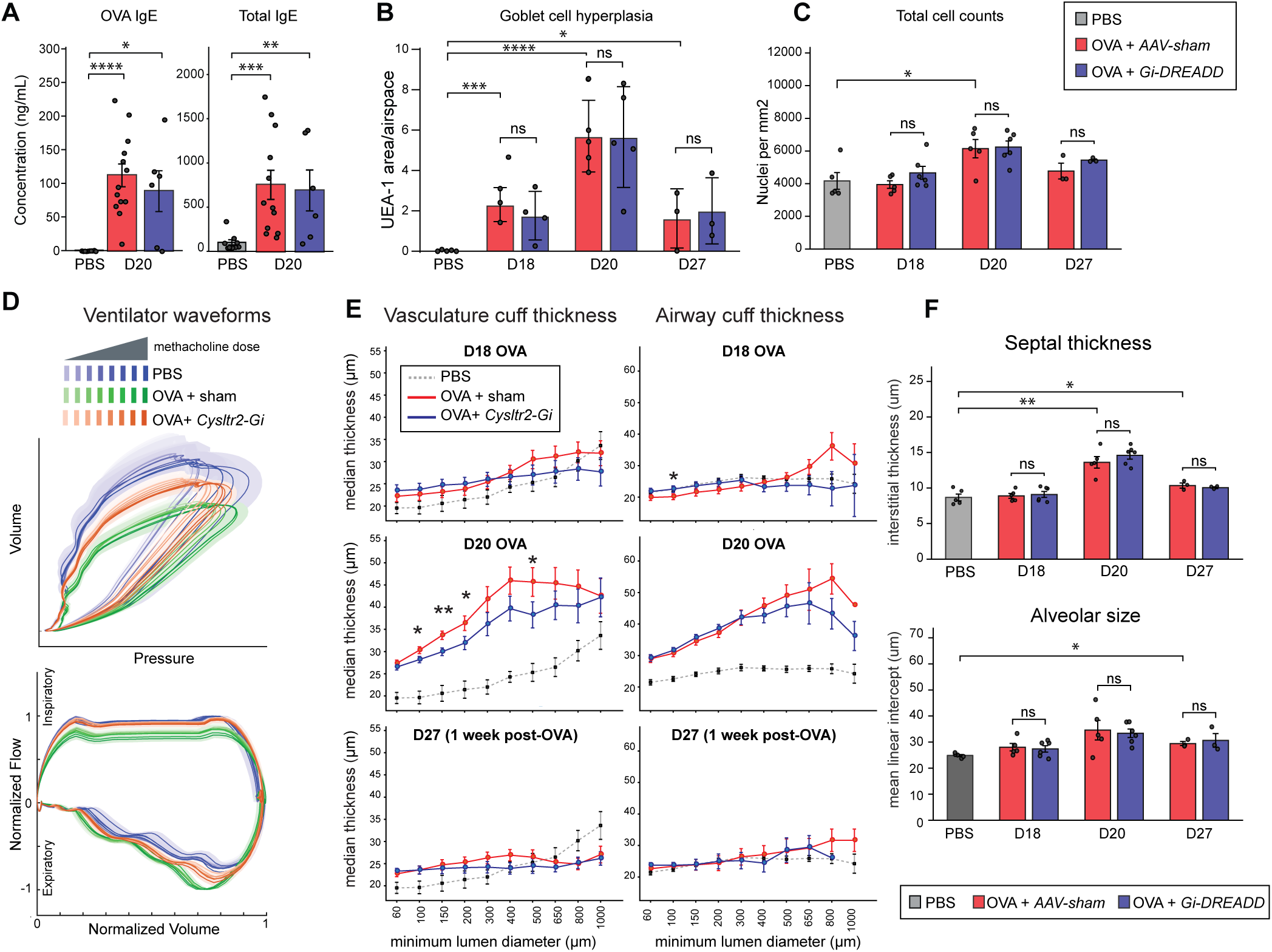
Silencing *Cysltr2*⁺ neurons rescues lung function and structure during severe allergic challenge. **(A)** OVA-specific and total IgE (μg/mL) in serum collected at day 20 from saline controls (gray), OVA-treated *Cysltr2-ires-Cre* mice injected in the nodose with *AAV-Flex-tdTomato* (pink), and OVA-treated *Cysltr2-ires-Cre* mice injected with *AAV-hSyn-DIO-hM3D(Gi)-mCherry* (blue). The same cohorts and colors apply to (B) to (F). n = 6-12 mice per group; Welch’s unpaired t test. **(B)** UEA-1⁺ area normalized to bronchial area in lung cryosections collected at days 18, 20, and 27 (4 to 6 lobes per mouse; n = 3–5 mice per group); two-tailed unpaired t test. **(C)** Total nuclei counts (nuclei per mm^2^) for the same cohorts at days 18, 20, and 27. n = 3-6; Welch’s unpaired t test. (**D)** Pressure-volume loops for each cohort. **(E)** Extended vascular and airway cuff thickness at days 18, 20, and 27. n = 3-6; Welch’s unpaired t test. **(F)** (Top) Septal/Interstitial thickness from H&E sections at days 18, 20, and 27. (Bottom) Alveolar size measured using mean linear intercept (μm) from H&E sections at days 18, 20, and 27; lobes were inflation-normalized as described in the supplementary materials. n = 3-6; Welch’s unpaired t test. Asterisks as in Fig. S1E; data are means ± SEM.

**Fig. S6.**
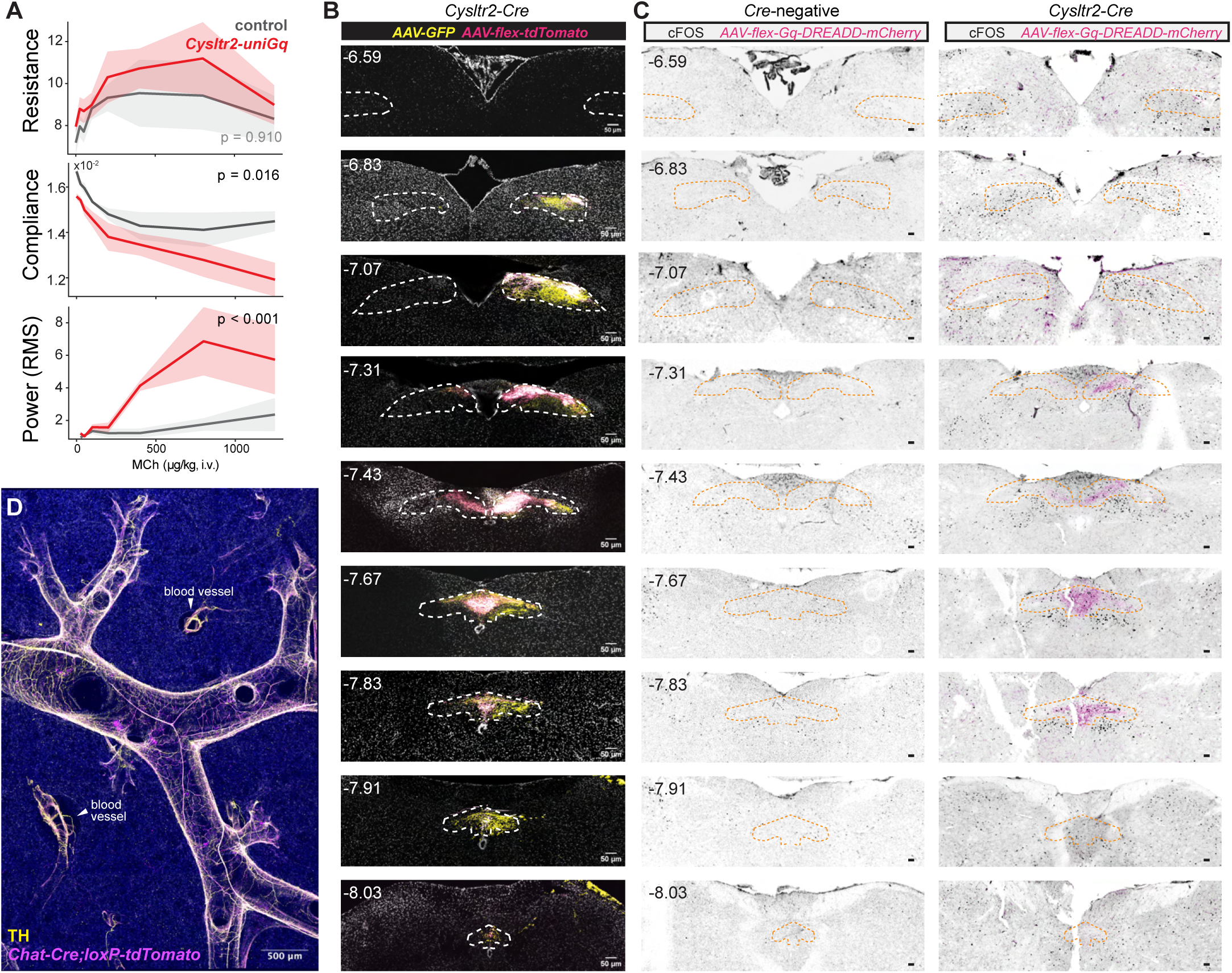
Characterization of neural pathways regulating pulmonary vascular fluid flux. **(A)** Resistance (AUC), compliance (median), and wave power during MCh challenge (i.v.; 25, 50, 100, 200, 400, 800, 1250 μg/kg) in *AAV-Flex-tdTomato*-injected controls (gray, n = 4) and *Cysltr2-ires-Cre* mice with unilateral *AAV-hSyn-DIO-hM3D(Gq)-mCherry* injection (red, n = 3). Omnibus model as in Fig. S1D; data are means ± SEM. **(B)** GFP and RFP immunostaining in *Cysltr2-ires-Cre* mice unilaterally injected with *pENN.AAV.CB7.CI.eGFP.WPRE.rBG* and *AAV-Flex-tdTomato*. **(C)** c-FOS immunostaining in serial brainstem sections 90 minutes after C21 in *Cre*-negative controls (left) and *Cysltr2-ires-Cre* mice (right). c-FOS⁺ cells were enriched in the m-NTS from bregma −7.31 to −7.43 mm. Outlines delineate NTS subregions at the indicated bregma levels. Scale bars, 50 µm. **(D)** Top: TH (yellow) and CHAT (magenta) immunostaining in a whole-mount lung slice (500 μm); nuclei, DAPI (blue). Blood vessels indicated on image. Scale bar, 500 μm.

**Fig. S7.**
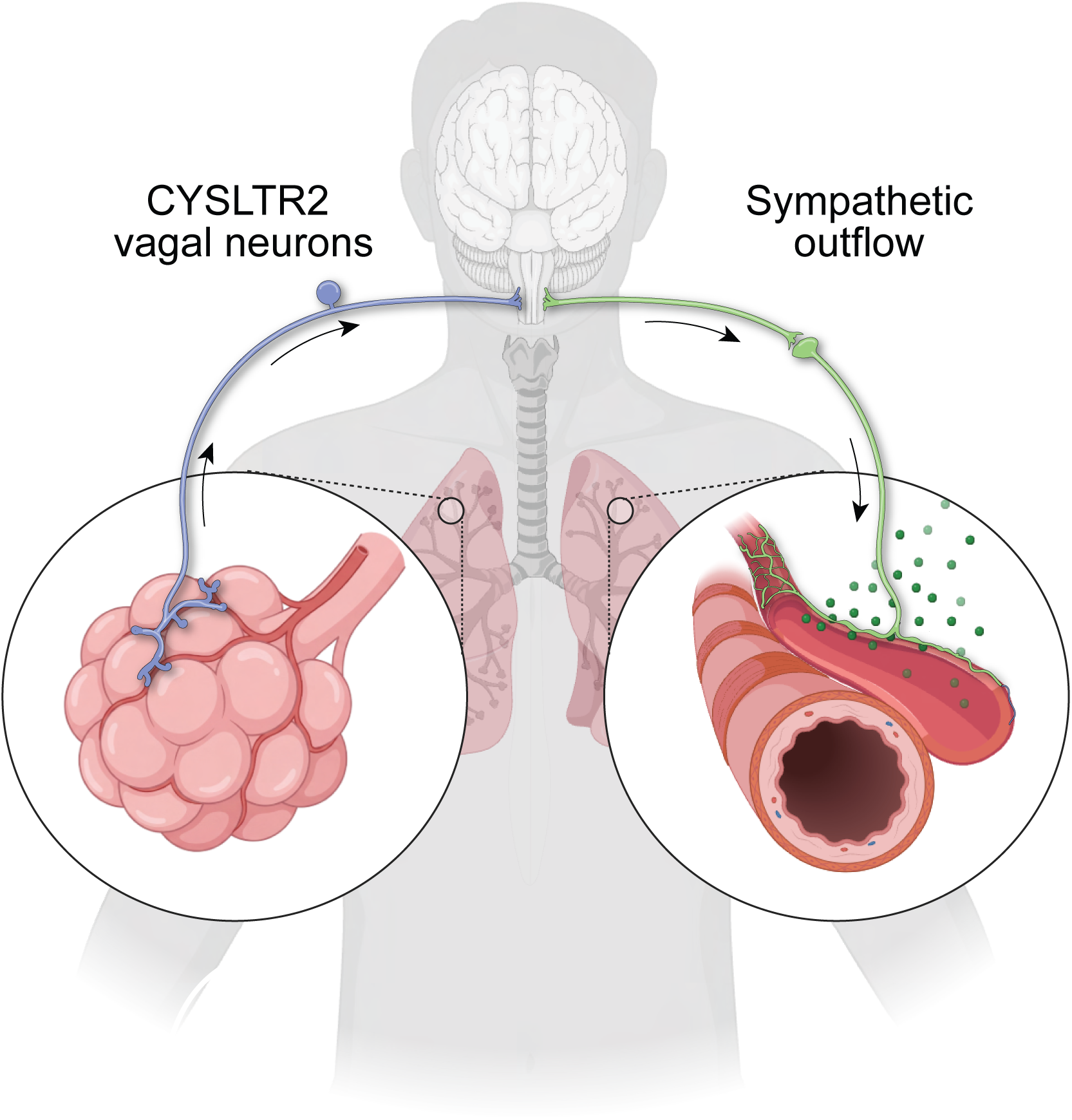
Proposed circuit by which CYSLTR2⁺ vagal sensory neurons drive sympathetically mediated pulmonary vascular leak.

**Table S1.**
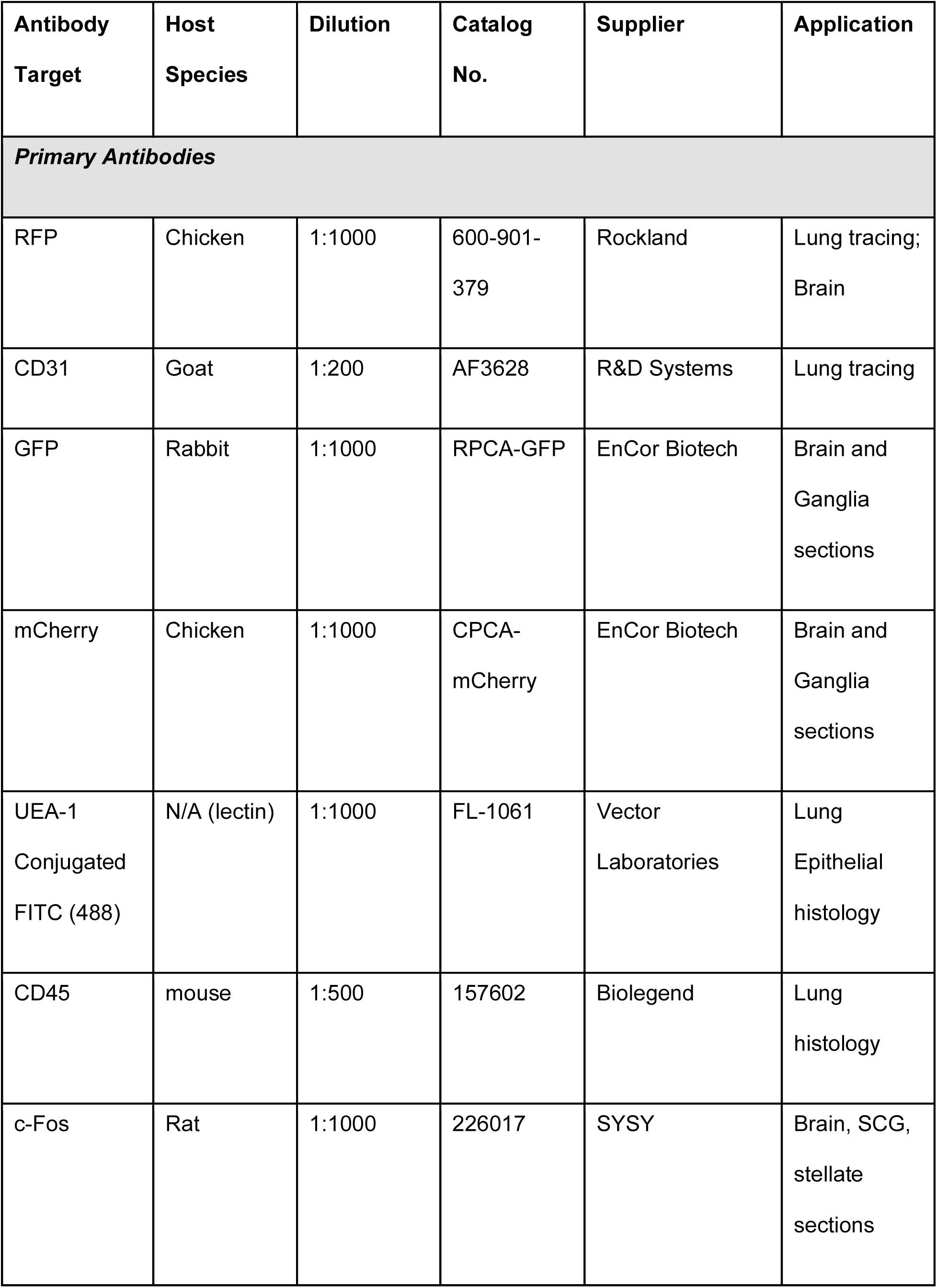
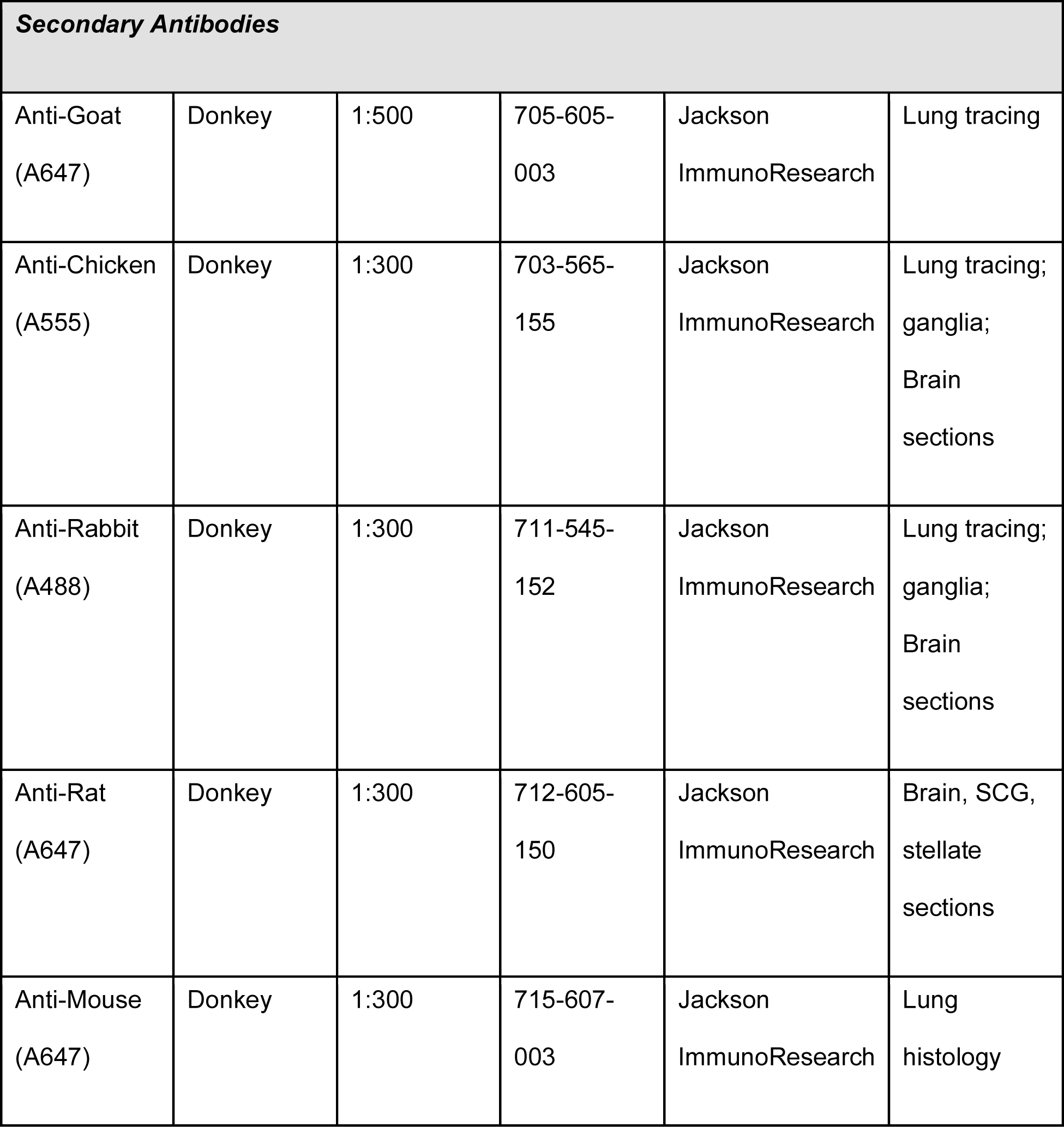
Primary and Secondary Antibodies for Immunostaining. *A = Alexa Fluor dye conjugate*.

| Antibody<br>Target | Host<br>Species | Dilution | Catalog<br>No. | Supplier | Application |
| --- | --- | --- | --- | --- | --- |
| **Primary Antibodies** |  |  |  |  |  |
| RFP | Chicken | 1:1000 | 600-901-379 | Rockland | Lung tracing;<br>Brain |
| CD31 | Goat | 1:200 | AF3628 | R&D Systems | Lung tracing |
| GFP | Rabbit | 1:1000 | RPCA-GFP | EnCor Biotech | Brain and<br>Ganglia<br>sections |
| mCherry | Chicken | 1:1000 | CPCA-mCherry | EnCor Biotech | Brain and<br>Ganglia<br>sections |
| UEA-1<br>Conjugated<br>FITC (488) | N/A (lectin) | 1:1000 | FL-1061 | Vector<br>Laboratories | Lung<br>Epithelial<br>histology |
| CD45 | mouse | 1:500 | 157602 | Biolegend | Lung<br>histology |
| c-Fos | Rat | 1:1000 | 226017 | SYSY | Brain, SCG,<br>stellate<br>sections |

| **Secondary Antibodies** |  |  |  |  |  |
| --- | --- | --- | --- | --- | --- |
| Anti-Goat<br>(A647) | Donkey | 1:500 | 705-605-003 | Jackson<br>ImmunoResearch | Lung tracing |
| Anti-Chicken<br>(A555) | Donkey | 1:300 | 703-565-155 | Jackson<br>ImmunoResearch | Lung tracing;<br>ganglia;<br>Brain<br>sections |
| Anti-Rabbit<br>(A488) | Donkey | 1:300 | 711-545-152 | Jackson<br>ImmunoResearch | Lung tracing;<br>ganglia;<br>Brain<br>sections |
| Anti-Rat<br>(A647) | Donkey | 1:300 | 712-605-150 | Jackson<br>ImmunoResearch | Brain, SCG,<br>stellate<br>sections |
| Anti-Mouse<br>(A647) | Donkey | 1:300 | 715-607-003 | Jackson<br>ImmunoResearch | Lung<br>histology |

**Table S2:** Color Deconvolution Settings.

|  |  |  |
| --- | --- | --- |
| **Color 1 (Evans**<br>**Blue)** | **Color 2 (red-**<br>**blood)** | **Color 3**<br>**(green)** |
| 0.69890785 | 0.37861538 | 0.41732463 |
| 0.64401823 | 0.7191889 | 0.7968193 |
| 0.31107616 | 0.58259565 | 0.43694293 |

**Table S3:** Combined GSEA across all vagal subtypes. Significant gene set enrichment analysis (GSEA) results for N10 and N12, analyzed across all vagal subtypes.

**Movie S1. Three-dimensional reconstruction of CYSLTR2+ vagal sensory neuron projections in the pulmonary alveolar parenchyma.** Three-dimensional rendering of virally labeled CYSLTR2+ vagal sensory neuron fibers (white) extending around a major conducting airway labeled with smooth muscle actin (green) and projecting into the alveolar parenchyma. CD31+ blood capillaries are labeled in purple, and nuclei are labeled with DAPI (blue).

**Movie S2. High magnification three-dimensional reconstruction of CYSLTR2+ vagal sensory neuron projections in the alveolar parenchyma.** Three-dimensional rendering of virally labeled CYSLTR2+ vagal sensory neuron fibers (white) innervating the alveolar parenchyma, with CD31+ blood capillaries labeled in purple and nuclei labeled with DAPI (blue). The imaged region corresponds to that shown in Fig. 2B.

## Notes

### Competing Interest Statement

The authors have declared no competing interest.

